# Preclinical evaluation of Targeted IL-1β Knockdown via CD44-Immunoliposomes: A Nano-therapy against the Inflammatory Microenvironment

**DOI:** 10.1101/2025.10.17.683015

**Authors:** Haly Shukla, Simran Nasra, Milonee Patel, Dhiraj Bhatia, Ashutosh Kumar

**Affiliations:** Biological and Life Sciences, School of Arts & Sciences, Ahmedabad University, Central Campus, Navrangpura, Ahmedabad 380009, Gujarat, India; Department of Biological Sciences and Engineering, Indian Institute of Technology Gandhinagar, Palaj, Gandhinagar – 382055, Gujarat, India

**Author notes:** These authors contributed equally to this work (Haly Shukla, Simran Nasra). Corresponding Author: Ashutosh Kumar, Associate Professor, Biological & Life Sciences, School of Arts & Sciences, Ahmedabad University, Central Campus, Navrangpura, Ahmedabad 380009 Gujarat, India.

**Keywords:** RNA therapeutics, IL-1β, Immunoliposomes, Macrophages, T-cells and Inflammation

## Abstract

Chronic inflammation, characterized by the infiltration of macrophages and the heightened release of pro-inflammatory cytokines, is the underlying cause of the pathogenesis of many critical diseases. Therapeutic interventions for controlling inflammation via gene knockdown of inflammatory mediators have emerged as a promising approach for regulating uncontrolled inflammation. This study explores the potential of siIL-1β-anti-CD44-Liposomes (SIL) as a potent anti-inflammatory therapy against pro-inflammatory RAW264.7 macrophages via gene specific knockdown of IL-1β mRNA through RNAi, and the subsequent down-regulation of the pro-inflammatory cytokine loop. The designed SIL exhibited a uniform size of 131.1 ± 0.5 nm with a quasi-spherical morphology and sustained release of siIL-1β within 24 hours. The reduction in pro-inflammatory cytokines like IL-1β, TNF-α, and IL-6 and inflammatory enzymes iNOS and COX-2; and the simultaneous increase in the anti-inflammatory cytokine IL-4, is indicative of the formulation’s therapeutic efficacy in reducing inflammation at a cellular level. The effects of SIL on the Macrophage-T cell crosstalk also uncovers the liposome’s efficacy in reducing cytokine-mediated T cell effector functions. The nuanced effects of siIL-1β-anti-CD44-Liposomes on *in-vivo* model of chronic inflammation underscore their potential for precise therapeutic interventions in inflammatory conditions, with multifaceted anti-inflammatory effects on tissue levels and cytokine levels.

Schematic representation of the study: CD44 receptors are elevated in LPS activated macrophages, which can be targeted by using anti-CD44 Liposomes, for RNA therapy. IL-1β knockdown via siRNA leads to lower inflammatory nature of macrophages, compromising it’s the antigen presentation and T-cell activation. This lowers the cytokine storm in inflammatory milieu and lower tissue damage can be achieved. Created with BioRender.com.

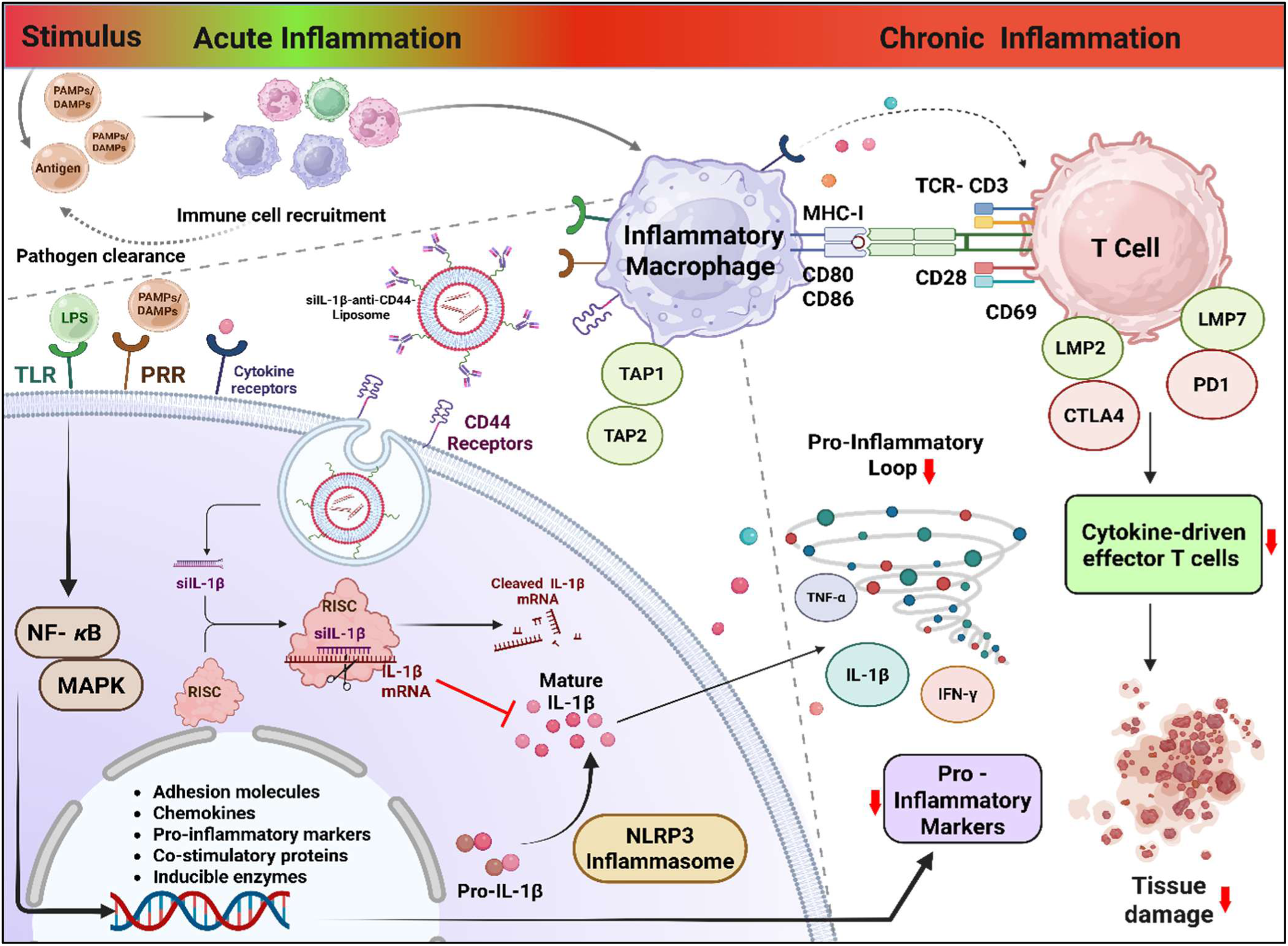

## 1. Introduction

Chronic inflammation plays a pivotal role in the pathogenesis of various diseases, including autoimmune disorders, and auto-inflammatory diseases, acting both as a contributing factor and a manifestation of these conditions. Tissue-specific inflammatory response leads to elevated levels of cytokines, adhesion molecules, and matrix metalloproteinase (MMPs) in the bloodstream; consequently resulting in the amplification of the inflammatory cascade^1^. Characterized by redness, pain, and swelling, uncontrolled inflammation can result in tissue damage and lead to organ dysfunction.

The key players of an inflammatory cascade are macrophages that function in phagocytosis, immunomodulation, and antigen presentation to the T cells^2^. Macrophages release certain inflammatory mediators like inducible nitric oxide (iNOS), which causes oxidative stress and tissue damage through nitric oxide release, cyclooxygenase (COX)-2, involved in the production of pain mediating prostaglandins, and inflammatory cytokines ^3^. The imbalance between pro-inflammatory and anti-inflammatory cytokines is a characteristic feature of refractory or severe chronic inflammatory conditions. Therapeutic interventions against these pro-inflammatory phenotypes of macrophages hold a lot of potential in controlling inflammatory conditions. These interventions focus on Macrophage Polarization from their pro-inflammatory (M1) phenotype to anti-inflammatory and pro-repair (M2) phenotype, reducing the levels of inflammation mediators like cytokines and adhesion molecules or inducing apoptosis in macrophages^4^. These interventions are often brought about using specific classes of drugs such as DMARDs, NSAIDs, and Biologics ^5^. While these therapies offer various mechanisms of controlling inflammation, their adverse side effects due to low bioavailability and non-specific activity render them insufficient. To address these concerns, nanotechnology has been used for targeted delivery of anti-inflammatory cargo as it offers better bioavailability, controlled release, and site-specific delivery of drugs to the desired cells^6^. Various nano-delivery systems like stealth nanoparticles, lipidoid nanoparticles, and polymerosomes have been used for targeting macrophages^7^. Among the various options, lipid-based nanoparticles, specifically nanoliposomes, stand out due to their biocompatibility, the ability to provide sustained release of cargo and the potential avenue of targeted delivery to specific cells by surface functionalization with antibodies and protein ligands ^8^. A study by Chiong et al. demonstrated that the liposomal delivery of piroxicam enhanced the drug’s anti-inflammatory properties on RAW264.7 *in-vitro* model ^9^. In a similar liposome-based study, Nasra et al. reported the use of the liposomal delivery system for MTX and RELA-siRNA targeting the Folate receptors over-expressed in pro-inflammatory macrophages as a potent anti-inflammatory therapy against RA ^10^. The study revealed the higher internalization of Folate-targeted liposomes compared to their non-targeted counterparts and the combinational effect of the drug and siRNA in causing NF-κB knockdown, underscoring the liposome formulation’s potential as a targeted delivery system.

Moreover, functionalizing the liposomes with receptors or antibodies offers great potential for targeting cell-specific antigens for precision therapy, via liposome opsonization and receptor-mediated endocytosis ^11^. For instance, in pro-inflammatory macrophages, CD44 receptors get overexpressed on the Macrophage cell surface in response to the up-regulated inflammatory cytokines TNF-α and IL-1β. This property of an inflammatory macrophage can be harnessed for targeted delivery via designing CD44-targeted liposome vehicles, as a potent anti-inflammatory therapy ^12^. These reports suggest the prospects of using liposomal delivery systems as a therapy against Macrophage-mediated inflammation. Among the various therapeutics delivered through liposomes, siRNA have emerged as promising candidates for interventions due to their crucial role in regulating gene expression by specific gene knockdown. Liposomal delivery facilitates the siRNA-mediated gene knockdown by protecting the siRNA from immune clearance and nuclease degradation, enabling successful delivery to the target cell. Gene knockdown of inflammation mediators that enhance and sustain the inflammatory cascade, such as the pro-inflammatory cytokine IL-1β, appear as great treatment modalities to control heightened inflammation and reduce the severity of damage caused by chronic inflammatory conditions. This study aims to harness the overexpression of CD44 receptors on an inflammatory macrophage for targeted delivery via nano-liposomes to deliver siRNA against the pro-inflammatory cytokine IL-1β as a potent anti-inflammatory therapy. The designed siIL-1β-anti-CD44-Liposomes (SIL) exhibited a uniform size of 131.1 ± 0.5 nm with high stability (over 10 months). After confirmation of siRNA entrapment, the release profile was performed and the liposomes were tested for their anti-inflammatory effects. The anti-inflammatory properties of the liposomes were also studied on an LPS-induced *in-vivo* model of chronic inflammation.

## 2. Results and Discussion

### 2.1 Formulation and characterization of siIL-1β encapsulated liposomes functionalized with CD44 Monoclonal Antibody (siIL-1β-anti-CD44-Liposomes)

The nanoparticle size and surface charges are pivotal measures in efficient design of stable delivery systems for siRNA. A noticeable increase was observed in the size of the anti-CD44-Liposomes (146.06 ± 1.52 nm) as compared to the empty liposome formulation (127.6 ± 4.6nm). This increase in size maybe an effect of surface modification of the liposome due to conjugation with anti-CD44 monoclonal antibody (mAb). Moreover, the ζ-potential of the anti-CD44-Liposomes also reduced to −8.13mV compared to empty liposomes (1.1mV). A near-neutral value of ζ -potential has the tendency to form aggregates, while a higher magnitude of ζ -potential assures steric hindrance between the liposome entities implying better colloidal stability and increased shelf life ^13^. This suggests the high stability of anti-CD44-Liposomes compared to empty liposomes and their enhanced potential to be efficient vehicles for siRNA delivery.

The hydrodynamic size of the siIL-1β-anti-CD44-Liposomes was estimated to be 131.1 ± 0.5 nm, with a ζ -potential of −6.73 mV. This slight decrease in the size of the anti-CD44-Liposomes upon siRNA loading might be due to compression due to the siRNA’s interaction with the lipid bilayer via electrostatic attractions between negatively charged siRNA backbone and positively charged lipid moieties. Moreover, the higher negative ζ -potential underscores the high stability of the liposome formulation. Additionally, the liposome formulation exhibited a very low Polydispersity Index (PDI) value of 0.073, indicating a narrow size distribution which implies that all particles had uniform size; suggesting the homogeneity of the final siIL-1β-anti-CD44-Liposome formulation. As indicated in Figure 1 a-b, d-e, the Hydrodynamic size and ζ-Potential of all liposome formulations were preserved over the span of 30 days. The siIL-1β-anti-CD44-Liposomes exhibited exceptional stability compared to its vehicle control counterparts, maintaining consistent size and PDI even after 10 months, whenever randomly measured. This signifies their high suitability as delivery systems, ensuring their stability, and insinuating their potential success in therapeutic applications for biological systems. The size, PDI and ζ-Potential of different formulations are listed in Figure 1f.

**Figure 1.**
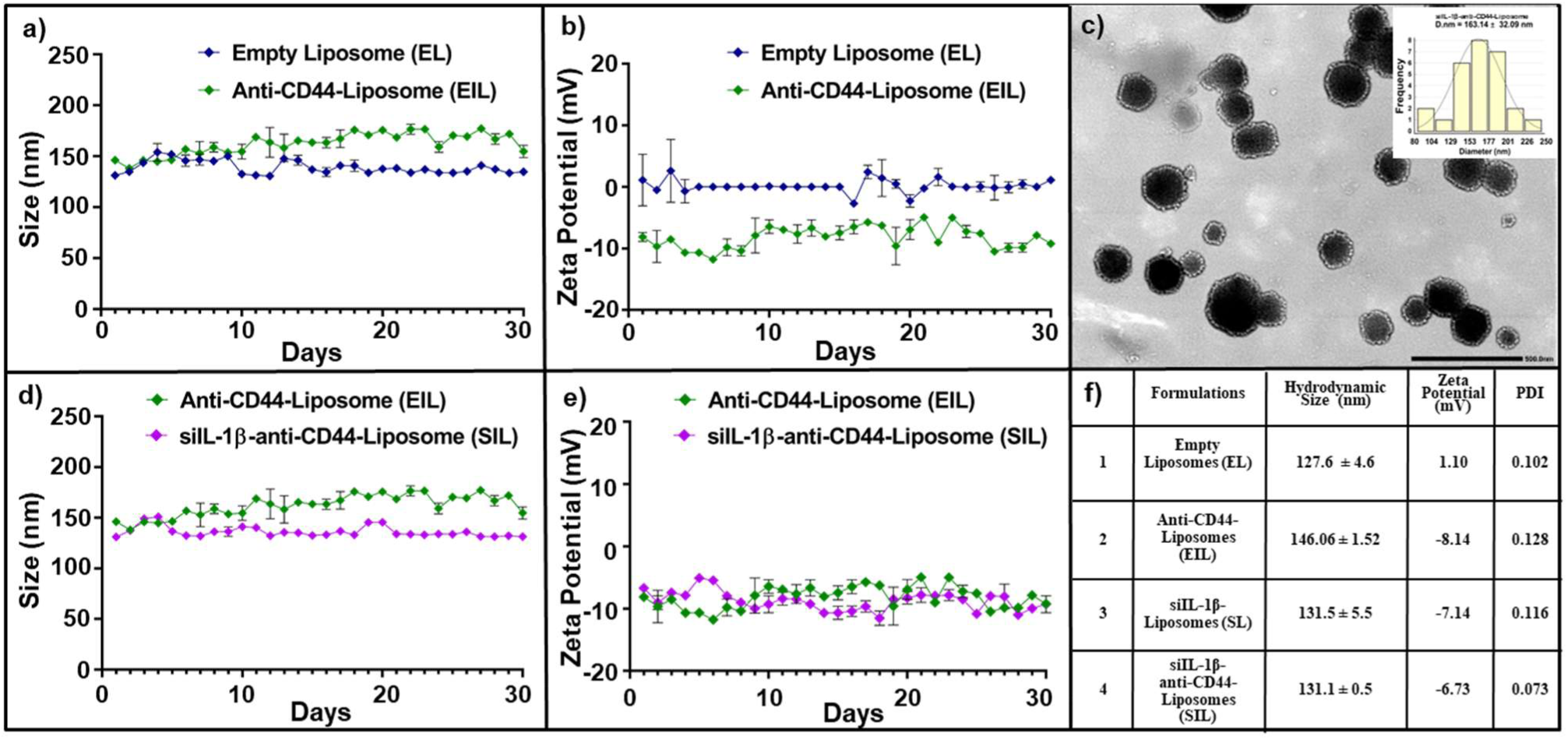
Formulation and characterization of liposomes- a) and b) represent the comparative stability in Hydrodynamic size and ζ-Potential of Empty Liposomes and anti-CD44-Liposomes over 30 days c) The quasi-spherical morphology of siIL-1β-anti-CD44-Liposomes and their size distribution with the average d.nm as 163.1 ± 32.09nm; 500nm x 500nm areas were visualized d) and e) depict the comparative stability in Hydrodynamic size and ζ-Potential of anti-CD44-Liposomes and siIL-1β anti-CD44-Liposome formulation over 30 days f) shows the compiled hydrodynamic size, ζ-Potential and PDI of all the liposome formulations on the day of preparation.

The morphological analysis of the liposomes was performed through TEM. As seen in Figure 1c, a quasi-spherical structure of liposomes was observed, and the siIL-1β-anti-CD44-Liposomes exhibited an average size of 163.1 ± 32.09 nm, which was remarkably high from the size of empty liposome formulation (96.2 ± 17.9 nm) (Refer-SI1). This size increase signifies the modifications in the liposome core followed by surface modification with antibodies.

#### 2.1.1 Validation of successful functionalization of CD44 mAb on the surface of Liposomes

Schematic representation of the study: CD44 receptors are elevated in LPS activated macrophages, which can be targeted by using anti-CD44 Liposomes, for RNA therapy. IL-1β knockdown via siRNA leads to lower inflammatory nature of macrophages, compromising it’s the antigen presentation and T-cell activation. This lowers the cytokine storm in inflammatory milieu and lower tissue damage can be achieved. Created with BioRender.com. Thiolation reaction involves the interaction between 2-iminothiolane (Traut’s Reagent) with the anti-CD44 mAb, through its amine group, leading to the addition of sulfhydryl residues on the mAb surface; as depicted in Figure 2a. After thiolation, the unbound anti-CD44 mAb was eliminated by affinity purification through the PD-10 desalting column. Figure 2c represents the concentration of elution fractions of anti-CD44 alone and thiolated anti-CD44 are displayed in blue and red regions, respectively. The reappearance of peaks corresponding to anti-CD44 mAb in the elution profile of the thiolated mAb, represented by the overlapping purple in fractions 3, 4 and 5, confirms thiolation. Therefore, the eluted Fractions 3, 4, 5 of the thiolated mAb were used for anti-CD44-Liposomes preparation owing to their high CD44 concentration. As observed in Figure 2b, a standard CD44 mAb has Amide-I and Amide-II bonds at the wavelength ranges of 1650–1655 cm^-1^ and 1540–1550 cm^-1^, respectively, and an additional N-H stretch at 3100-3600 cm^-1^. The reflection of these peaks in the thiolated mAb sample further confirms the presence of mAb in the eluted fractions, along with the appearance of an additional peak at 2576cm^-1^. This peak corresponds to the sulfhydryl bond, aligning with the reported range 2550–2600 cm^-1^, introduced after the reaction with Traut’s reagent, confirming the thiolation of anti-CD44 mAb.

**Figure 2.**
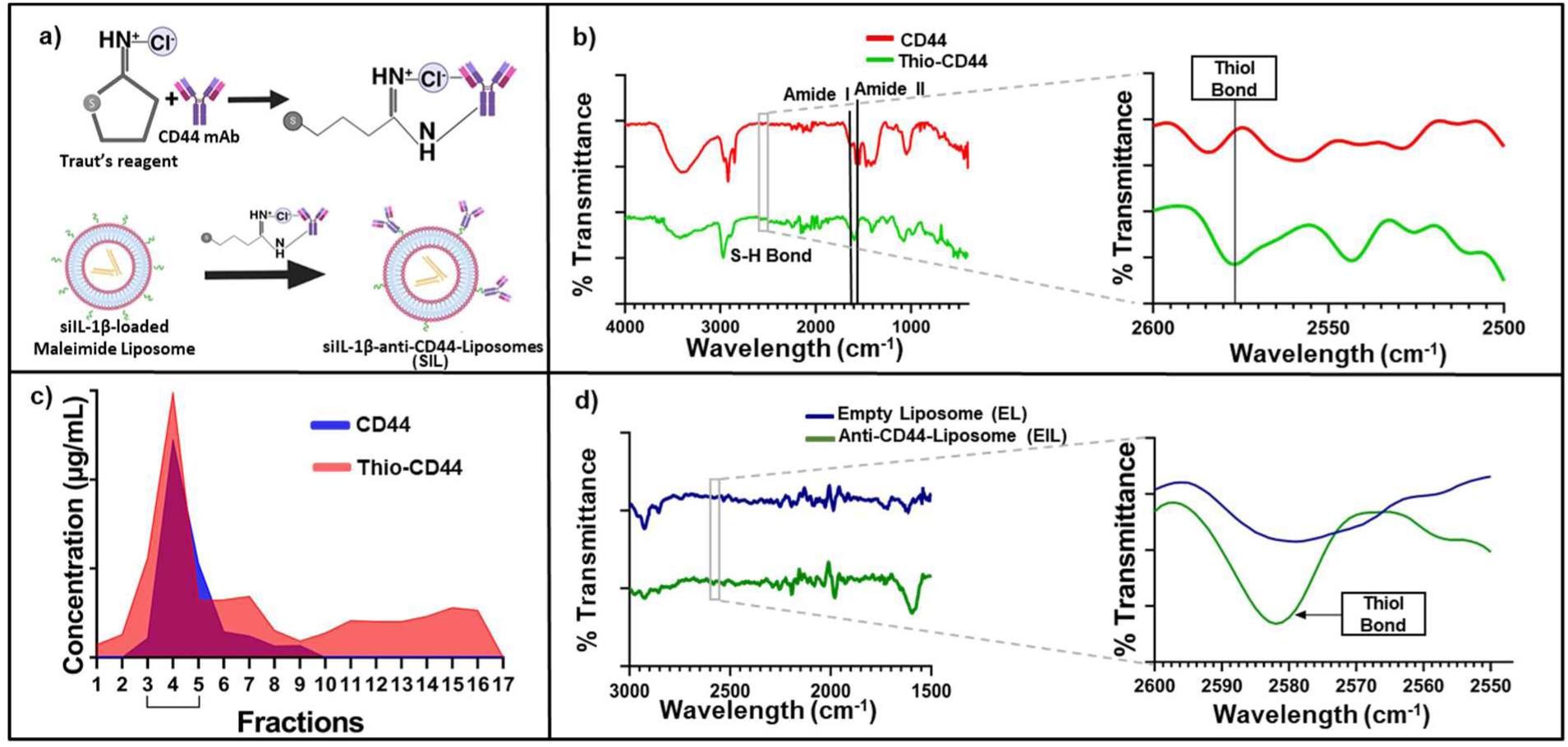
a) Schematic illustration of Thiolation reaction and attachment of thiolated mAb onto the surface of Maleimide Liposomes b) Confirmation of thiolation of anti-CD44 mAb using FTIR c) The Concentrations of anti-CD44 mAb and Thiolated anti-CD44 elution fractions. Fractions 3, 4 and 5 with highest concentration of anti-CD44 mAb were used for the preparation of anti-CD44-Liposomes d) Confirmation of mAb conjugation with malemide liposome through thiol (S-H) bond present in FTIR peak.

The conjugation of thiolated mAb on anti-CD44-Liposomes was validated by FTIR as reported in Figure 2d. The peaks of the amide bonds in the mAb were seen in the FTIR spectra of siIL-1β-anti-CD44-Liposomes, which were absent on the empty liposomes. The peaks corresponding to the thiols bond were observed in its ideal range of 2550–2600 cm^-1^, specifically at 2597 cm^-1^, further confirming the conjugation of mAb on the liposome surface.

### 2.2 Optimization of N/P Ratio, evaluation of the Encapsulation Efficiency and their Release Kinetics

The formation of siIL-1β-Liposomes takes place on an optimum ratio of the positive charges from lipids to negative charges from siRNA. This makes the N/P ratio a critical parameter in designing efficient delivery systems for siRNA. The N/P ratio, pivotal in determining the complexation efficiency of the siRNA in the liposomes, ensures encapsulation by considering the concentration of the Nitrogen from Lipids to the Phosphates from siRNA backbone. The gel retardation assay confirms the presence of unbound siRNA in different N/P ratios, and no bands upon successful encapsulation. As shown in Figure 3 a, c, the N/P ratio 8 was used for preparing the liposome formulation, as the gel retardation assay depicted the absence of siRNA bands after dialysis. The siIL-1β-Liposomes were found to have a very high encapsulation Efficiency close to 99% as depicted in Figure 3 d. The release kinetic profile of the siIL-1β-Liposomes showed an incremental release of the siRNA by 63% for the first 8 hours, followed by a sustained release till the 24^th^ hour, and an almost 100% release in 48 hours. The standard curve is presented in Figure 3b. The high entrapment efficiency and release profile underscores the formation of an effective delivery system for IL-1β siRNA for targeted gene knockdown.

**Figure 3.**
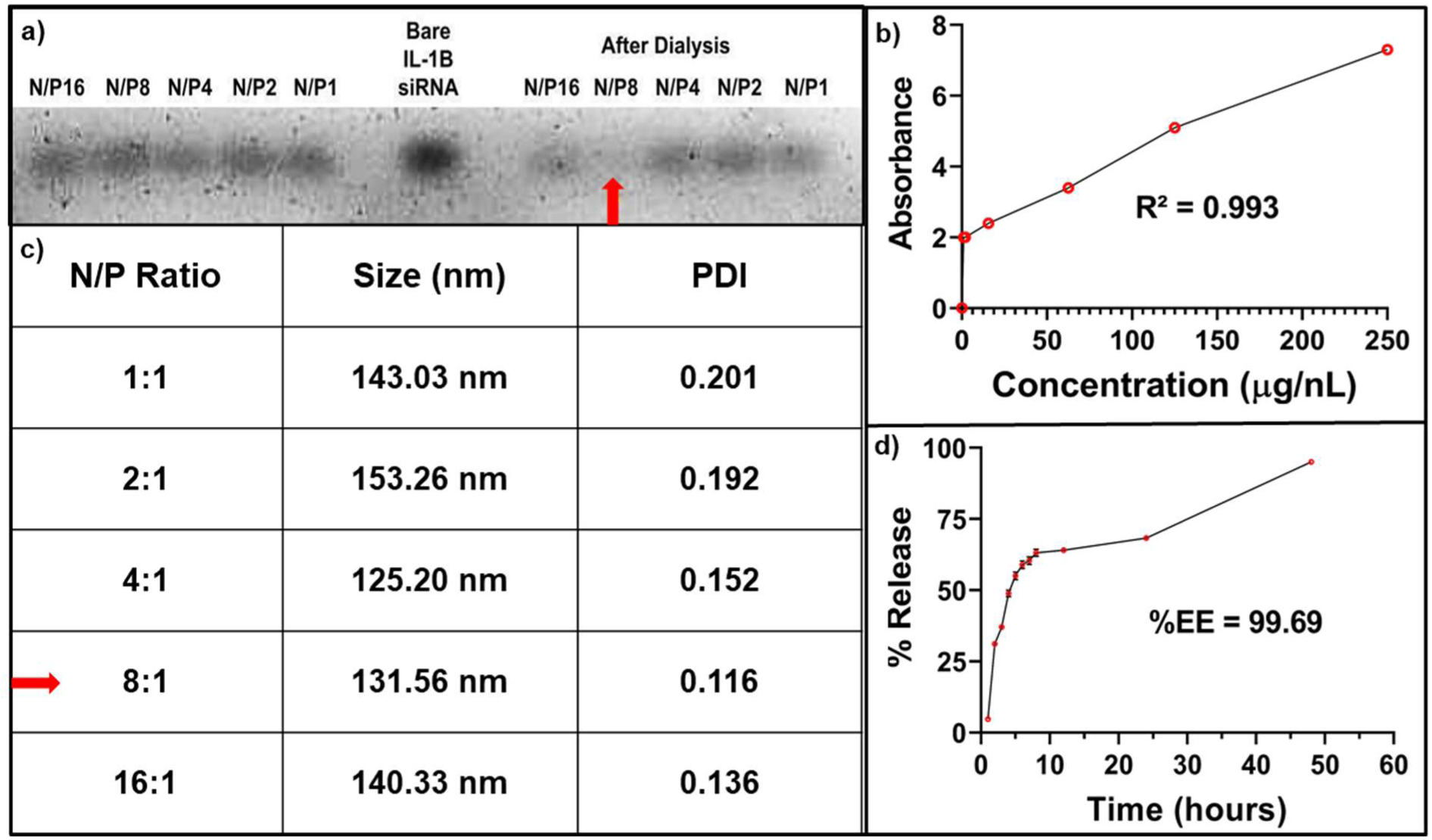
a) and c) Depicts the N/P ratio obtained at 8 in the Gel Retardation Assay and the size and PDI of the Liposome formulations at different N/P Ratios, respectively. b) The calibration curve of siIL-1β standards d) Release Profile of siIL-1β achieved in 48 hours, and the %EE obtained at 99.69%

### 2.3 Successful internalization of anti-CD44-Liposomes in activated RAW264.7 macrophages

The relative uptake of Rhodamine loaded Liposomes and Rhodamine loaded anti-CD44-Liposomes in RAW264.7 macrophage cell line was estimated through Confocal Microscope, at an objective of 100X. The naive M0 macrophages were activated to its pro-inflammatory phenotype using LPS to mimic the inflammatory tissue microenvironment. Naïve macrophage (referred as M0) exhibit a spherical morphology, which transforms to an elongated shape upon LPS stimulation, typically characterized by the presence of cytoplasmic extensions, as depicted in Figure 4a, with yellow arrows.

**Figure 4.**
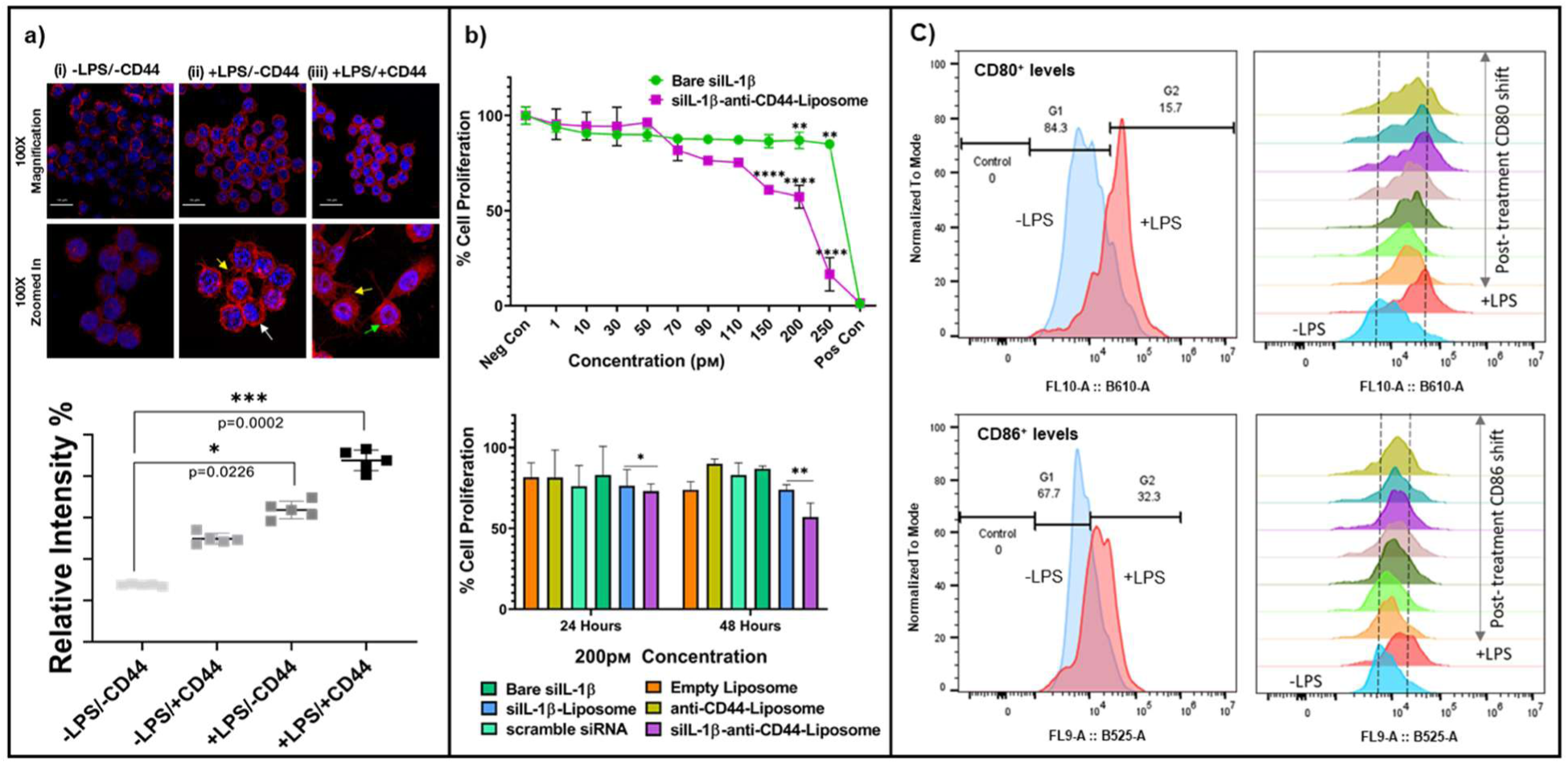
a) Pictorial and graphical representation of enhanced uptake of Rhodamine after anti-CD44 mAb functionalization on liposomes as observed through Confocal Microscopy, magnification 100X. The p-values were calculated through Kruskal–Wallis test b) Comparative analysis of cytotoxicity induced by siIL-1β-anti-CD44-Liposomes as compared to bare IL-1β; IC_50_ of siIL-1β-anti-CD44-Liposomes as 225.6 pM, after 48 hours b) The rate of cell proliferation under different treatment conditions, at 200pM. All experiments were performed in biological and technical triplicates. Two-way ANOVA was used to test significance c) CD80 and CD86 levels increase upon LPS activation and these levels gets down-regulated post treatment of siIL-1β-anti-CD44-Liposomes. Concentration dependent decrease in CD80 and CD86 levels were observed. Orange peak: 10 pM, light green: 25 pM, dark green: 50 pM, light purple: 100 pM, dark purple: 125 pM, teal: 150 pM, and dark yellow: 175 pM. The p-values for these graphs are represented as * for p < 0.05, ** for p < 0.01, and *** for p < 0.001.

As represented in Figure 4a (i), the naive Macrophages without any LPS treatment show a minimal uptake of Rhodamine Liposomes. However, the naive Macrophages targeted with Rhodamine loaded anti-CD44-Liposomes exhibited a relatively higher rhodamine uptake, suggesting higher internalization. Activated macrophages have higher expression of CD44 receptor on their cell surface for increased cellular adhesion, and immune cell-recruitment ^14^. This up-regulation of CD44 receptors in LPS-activated macrophages can be exploited for targeted drug delivery through receptor-mediated endocytosis. The rhodamine internalization increased 3-fold in LPS-activated macrophages when treated with rhodamine loaded anti-CD44-Liposomes (Figure 4a (iii)) in contrast to naive macrophages. This highlights the ability of the liposomes to internalize more in LPS-activated macrophages, when surface modified with anti-CD44 mAb.

Another notable observation was the localization of rhodamine uptake in the cytoplasm in LPS-activated macrophages treated with rhodamine loaded anti-CD44-Liposomes, indicated with green arrows in Figure 4a (iii). In contrast, the rhodamine uptake in LPS activated cells after treatment with Rhodamine Liposomes without CD44 mAb shows the dye uptake only at the periphery, indicated with white arrows in Figure 4a (ii). This underscores the potential of anti-CD44-Liposomes in site-specific delivery, facilitating higher internalization of therapeutic agents to inflammatory tissue enriched in pro-inflammatory macrophages.

### 2.4 Quantitative assessment of cellular viability in RAW264.7 macrophages

The viability assay sheds light on the cytotoxicity generated by siIL-1β-anti-CD44-Liposomes on LPS-stimulated macrophages over duration of 24 hours and 48 hours. While administered alone, siIL-1β shows a negligible decrease in cell proliferation, with an estimated value of IC_50_ as 969 pM after 48 hours, upon extrapolation of the curve. In comparison to this, treatment with siIL-1β-anti-CD44-Liposomes exhibited a gradual decline in cell proliferation with an IC_50_ value of 225.6 pM after 48 hours (Figure 4b). This cytotoxicity is suggestive of the efficacy of siRNA to cause RNA interference; as minimal dosing of siRNA results in down-regulation of the target gene, but at higher levels, siRNA interferes with the cellular processes and disrupts them due to off-target gene silencing. This makes the IC_50_ a crucial determinant of the dosage at which gene silencing can be achieved using siRNA therapeutics. The dosage reduction of the siIL-1β-anti-CD44-Liposomes formulation compared to its vehicle control counterparts and bare IL-1β siRNA (Figure 4c) is also indicative of targeted delivery achieved through CD44, underscoring its therapeutic potential in IL-1β knockdown in LPS-activated macrophages through precise drug delivery.

### 2.5 Assessment of Co-stimulatory marker: CD80 and CD86 cell surface markers profiling

In macrophages, CD86 is typically expressed at higher levels compared to CD80, meaning resting macrophages (M0) primarily express CD86 while CD80 expression is usually low or absent unless stimulated by an inflammatory signal; both molecules are considered co-stimulatory markers important for T cell activation when expressed on antigen-presenting cells (APCs) like macrophages. As observed in Figure 4c, CD80 and CD86 levels increase after LPS activation in macrophages (red peak), indicating invasion of activated macrophages. Treatment concentration: 50 pM show most significant down-regulation of CD80 and CD86 levels that were elevated after LPS stimulation. IL-1β is a key driver of macrophage activation and promotes the expression of co-stimulatory molecules like CD80 and CD86, which are crucial for antigen presentation and T-cell activation. Lower IL-1β levels may reduce AP-1 and IRF-mediated transcriptional activation of CD80/CD86, impairing the macrophage’s antigen-presenting capacity and subsequently down-regulating the T-cell activation. The lower T-cell activation will decrease the further immune cell recruitment by dampening the cytokine storm.

### 2.6 Morphological changes upon LPS stimulation and post si-IL-1β-anti-CD44-Liposomes treatment

Figure 5a displays the changes observed in RAW264.7 macrophage cell line after LPS stimulation and further treatment with siIL-1β-anti-CD44-Liposomes. Unstimulated M0 macrophages appear as spherical shaped in SEM. Upon LPS activation, the cells displayed a larger elongated or flattened amoeboid morphology. The cells also exhibited cytoplasmic extensions and pseudopods, indicative of their enhanced phagocytic and migration abilities compared to their M0 counterparts ^15^. The observed changes in the morphology of cells after LPS stimulation are suggestive of their polarization towards the M1 phenotype ^16^. The M1 cells also display extracellular web-like structures, indicated with yellow arrows in Figure 4a, which may represent Macrophage extracellular traps (METs) ^17^. METs are a hallmark of immune defence mechanisms associated with tissue damage, heightened inflammation and autoimmunity. After treatment with siIL-1β-anti-CD44-Liposomes, the cells show a reduction in cytoplasmic extensions and METs indicating their shift towards a less inflammatory phenotype. The restoration of their small and spherical morphology depicts a distinctive feature of M2 macrophages, particularly the M2c phenotype associated with tissue repair ^18^.

**Figure 5.**
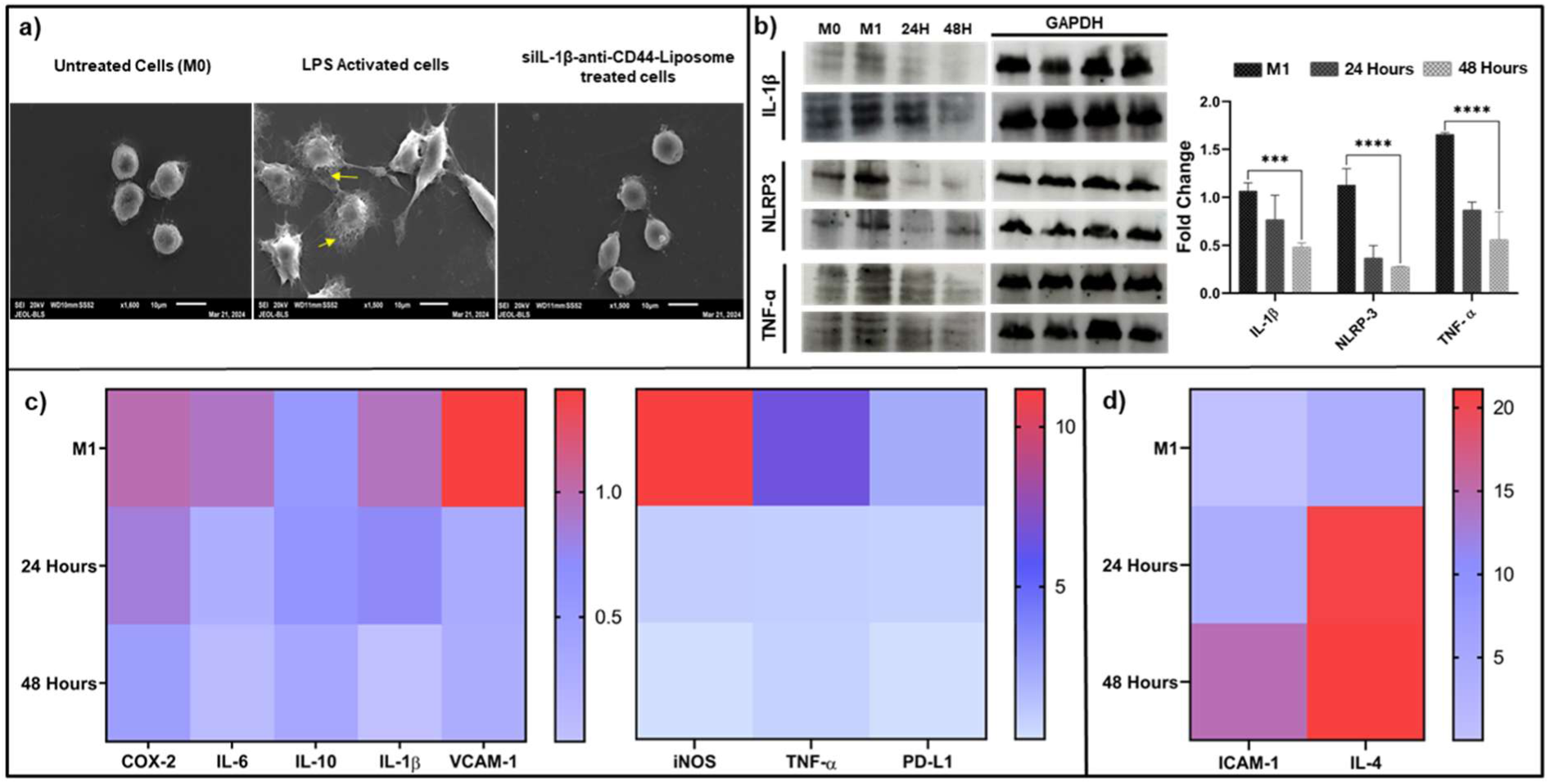
a) Morphological changes observed in RAW264.7 using Scanning Electron Microscope; 10µm x 10µm areas were visualized b) Western Blot analysis of key inflammatory proteins after LPS induction (represented as M1) and subsequent treatment with 150 pM siIL-1β-anti-CD44-Liposomes for 24 Hours and 48 Hours. The p-values for these graphs are represented as * for p < 0.05, ** for p < 0.01, *** for p < 0.001, and **** for p < 0.0001. Mean ± SD was plotted, and student’s t-test was used to determine significance. Analysis of gene expression on RAW264.7 cells c) down-regulated genes after 24 and 48 hours of siIL-1β-anti-CD44-Liposome treatment at 150pM and d) up-regulated genes after 24 and 48 hours of siIL-1β-anti-CD44-Liposome treatment at 150pM. The experiments were conducted in biological triplicates. Student’s t-test was used to determine significance

### 2.7 Quantitative gene expression and protein studies

Figure 5c displays the changes observed in the gene expression levels in the LPS-activated macrophages after siIL-1β-anti-CD44-Liposomes treatment. The down-regulation of the pro-inflammatory markers upon the liposome treatment sheds light on the anti-inflammatory properties of the therapy. The IL-1β knockdown after 48 hours post liposome treatment is indicative of the efficacy of both siRNA-delivery and the subsequent RNA interference pathway. The decrease in mRNA levels also correlated with the decrease in protein levels of IL-1β after siIL-1β-anti-CD44-Liposomes treatment, as depicted in Figure 5b. Moreover, the reduction in NLRP3 protein levels after siIL-1β-anti-CD44-Liposomes treatment highlights the liposome’s effectiveness in dampening the upstream pathway of IL-1β release.

Notably, a substantial down-regulation of the pro-inflammatory cytokine TNF-α was also observed after siIL-1β-anti-CD44-Liposomes treatment. This observation was also reflected at the protein levels with a significant reduction in TNF-α protein after 24 and 48 hours of siIL-1β-anti-CD44-Liposomes treatment (Figure 5b). TNF-α is an integral contributor of the inflammatory response as its downstream signalling pathway leads to the release of mitogen-activated protein kinases (MAPKs) and further activation of the NF-*κ*B signalling, which further elevates the levels of inflammation ^19^. The prominent feature of diseases driven by inflammation are the cytokine storms in their blood streams predominantly IL-1β, TNF-α and IL-6. The profound reduction of these pro-inflammatory cytokines after siIL-1β-anti-CD44-Liposomes treatment was indicative of the therapeutics’ effectiveness to attenuate inflammation on a broader level.

Another notable observation in gene expression levels was the sharp decline by 11-fold in the mRNA levels of iNOS after siIL-1β-anti-CD44-Liposomes treatment. iNOS is a crucial regulator of the Nitric Oxide synthesis in inflammatory conditions, and it’s up-regulation can cause severe damage to the underlying tissues. The release and over-expression of iNOS is heavily dependent on the environmental cues in the tissue microenvironment, pro-inflammatory cytokines being one of them ^20^. This reiterates the effectiveness of siIL-1β-anti-CD44-Liposomes in alleviating iNOS release through the knockdown of IL-1β and other pro-inflammatory cytokines, further underscoring its anti-inflammatory properties. Additionally, COX-2, a mediator of prostaglandin release responsible for redness, swelling and pain due to immune cell recruitment ^21^, was also significantly down-regulated upon siIL-1β-anti-CD44-Liposomes treatment. The reduction in the gene expression levels of PD-L1 further reaffirms these findings.

A further significant observation was the reduction in the levels of VCAM-1 which is responsible for immune cell adhesion and trans-endothelial migration during an inflammatory response. However, there were some noteworthy observations in the levels of ICAM-1 which had an increased expression after siIL-1β-anti-CD44-Liposomes treatment, as depicted in Figure 5d. Ideally, ICAM-1 is believed to contribute to enhanced adhesion between various cell types during inflammatory conditions but the increased levels of ICAM-1 after siIL-1β-anti-CD44-Liposomes treatment suggests its role in resolution of Inflammation. ICAM-1 has a pleiotropic role in mediating inflammatory responses, by inducing Macrophage polarization towards a pro-repair phenotype, and thus contributing in tissue repair ^22^. Studies by Neda et al and Gu et al further validate these properties exhibited by ICAM-1, highlighting its role in pro-repair reprogramming of macrophages to enhance tissue healing and inflammation resolution ^23^. Moreover, ICAM-1 also facilitates the immunological synapse between Macrophages and T cells, and supports the functioning of Macrophages during Antigen presentation. The dynamics of VCAM-1 suppression and ICAM-1 up-regulation suggest a controlled immune response associated with reduced inflammation due to less recruitment of T cells at the site of inflammation, while preserving macrophage functionality.

The dramatic increase by 20-fold in the levels of IL-4, an anti-inflammatory cytokine, upon siIL-1β-anti-CD44-Liposomes administration is indicative of the liposome’s ability to foster a shift towards a more anti-inflammatory environment. IL-4 is responsible for the alternate activation of macrophages, promoting their transition from the LPS-activated pro-inflammatory phenotype to the anti-inflammatory M2 phenotype, which is crucial for tissue repair and the resolution of inflammation. No dramatic changes in the expression of IL-10 indicate that the anti-inflammatory state is being controlled. The changes in gene expressions after siIL-1β-anti-CD44-Liposomes administration on LPS-activated macrophages thus indicates an overall suppression of the inflammatory response, characterized with down regulation of pro-inflammatory markers and an increase in anti-inflammatory markers. This underscores the liposome’s potential in minimizing inflammation-associated tissue damage, and its effectiveness as a promising therapy for chronic inflammatory conditions.

### 2.8 Macrophage mediated influence on T-cells

The co-culture of Macrophages and T cells revealed some intricate properties of immune response after treatment with siIL-1β-anti-CD44-Liposomes (Figure 6). An ideal inflammatory response is driven by an antigen or stimulus, which facilitates the formation of an immunological synapse between APCs (macrophages, dendritic cells and B cells) and T cells, further elevating the levels of inflammation. The binding of the TCR-CD3 complex with the antigenic peptide-MHC complex from APCs initiates the T cell-APC interaction, along with co-stimulatory molecules like CD28 and local cytokines. Activation with TCR results in increased survival and proliferation of T cells promoting their effector functions. These effector functions of T cells result in tissue damage in the surrounding tissues during chronic inflammatory conditions.

**Figure 6.**
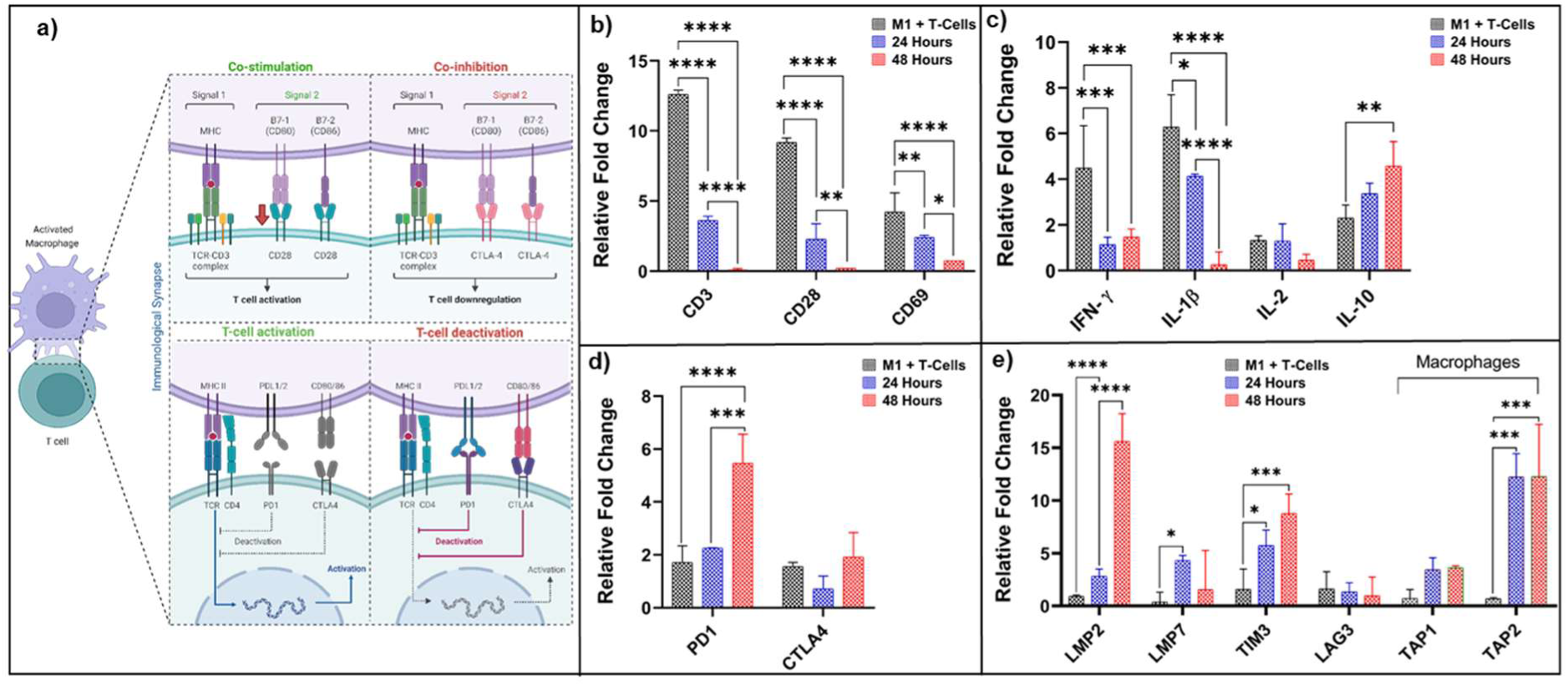
Gene expression levels observed after macrophage-T cell co-culture in-vitro, and the effect of siIL-1β-anti-CD44-Liposomes after 24 Hours and 48 Hours a) immune checkpoints b) CD markers of T cell activation c) cytokine levels d) regulatory proteins The p-values for these graphs are represented as * for p < 0.05, ** for p < 0.01, *** for p < 0.001, and **** for p < 0.0001. The experiments were conducted in biological triplicate, and a two-tailed t-test was used for statistical analysis.

A significant up-regulation in the Immune checkpoint PD-1 after treatment with siIL-1β-anti-CD44-Liposomes, along with a slight increase in the levels of CTLA-4, was observed, suggesting the inhibition of T cell activation. Both immune checkpoints are crucial markers of peripheral tolerance in T cells and help in regulating T cell functioning ^24^. CTLA-4 functions by producing inhibitory signals that act antagonistically to the co-stimulatory signals from the TCR-MHC complex. CTLA-4, along with CD28, the primary co-stimulatory marker in T cell activation, alternatively binds to the B7 ligands during T cell activation and inhibition, respectively ^25^. Thus, the up-regulation of CTLA-4 results in the direct inhibition of the co-stimulatory immune synapse due to disruption in CD28 binding. Moreover, CTLA-4 also enhances the mobility of T cells, thereby reducing the chances of interaction with APCs. PD-1 is also a key inhibitor of antigen receptor signalling, T cell proliferation, and subsequently contributes in the down-regulation of pro-inflammatory cytokines TNF-α and IFN-γ ^26^. Presence of PD-1 interrupts the phosphorylation of TCR signalling intermediates, thereby mitigating the early TCR signalling and consequently reducing the activation of T cells. This indicated the siIL-1β-anti-CD44-Liposomes’ efficacy in controlling the inflammation associated with T cell interactions with the APCs. This can be reaffirmed by the down-regulation in the gene expression levels of CD3, the main component of the TCR-APC interaction, and CD28, the co-stimulatory molecule that signals the priming of Naive T cells and promotes the TCR signalling. Additionally, siIL-1β-anti-CD44-Liposome treatment also attenuated the expression of CD69, an early activation marker responsible for T cell differentiation, which functions by elevating the levels of pro-inflammatory cytokines-TNF-α, IFN-γ and IL-2. CD69 is a prominent marker present in autoimmune conditions, and is associated with T cell-mediated tissue damage by promoting their effector functions ^27, 28^. Given these aspects of CD69 functioning, the effect of siIL-1β-anti-CD44-Liposomes on the CD69 levels is suggestive of the therapy’s potential in regulating inflammation. Moreover, a significant reduction in the levels of IFN-γ and IL-1β, pro-inflammatory cytokines, and a slight increase in IL-10, an anti-inflammatory cytokine were also observed.

A remarkable observation in the gene expression levels was the dramatic increase in the levels of LMP2 and a slight increase in LMP7; immune-proteasome units associated with antigen processing. Their up-regulation underscores their potential to establish immune homeostasis by preventing excess immunosuppression, a compensatory mechanism against the anti-inflammatory therapy ^29^. Moreover, LMP2 and LMP7 are also important for the resolution of inflammation as they facilitate antigen clearance and Immune adaptation. Additionally, an increase in the levels of TIM-3, a co-inhibitory molecule, also reiterates the deactivation of T cell effector functions in inflammation upon SIL treatment. No significant changes in the levels of LAG-3 were observed. TAP1 and TAP2 levels are up-regulated in macrophages, indicating better MHC-I response in macrophages. To summarize, the changes in gene expression levels of T cells after treatment with siIL-1β-anti-CD44-Liposomes reveal the anti-inflammatory and regulatory properties of the liposome formulation on antigen presentation and T cell effector functions. This property can be harnessed against T-cell-mediated diseases associated with chronic inflammation.

### 2.9 Assessment of Physiological Parameters During Treatment

To evaluate the systemic tolerability of different formulations, body temperature, food intake, and body weight were monitored throughout the experimental period (Figure 7). Body temperature remained stable within the physiological range across all groups, with only minor daily fluctuations. Food intake also remained largely consistent, indicating that none of the formulations adversely affected appetite or feeding behavior. Similarly, mean body weight trajectories demonstrated only modest variations across groups, with overlapping error margins, suggesting the absence of treatment-associated weight loss or systemic toxicity. These findings indicate that repeated administration of EIL, SL, or SIL did not result in overt physiological stress or toxicity in mice. The maintenance of body temperature, food intake, and body weight across groups supports the overall safety and tolerability of the formulations, which is crucial for their further preclinical evaluation.

**Figure 7.**
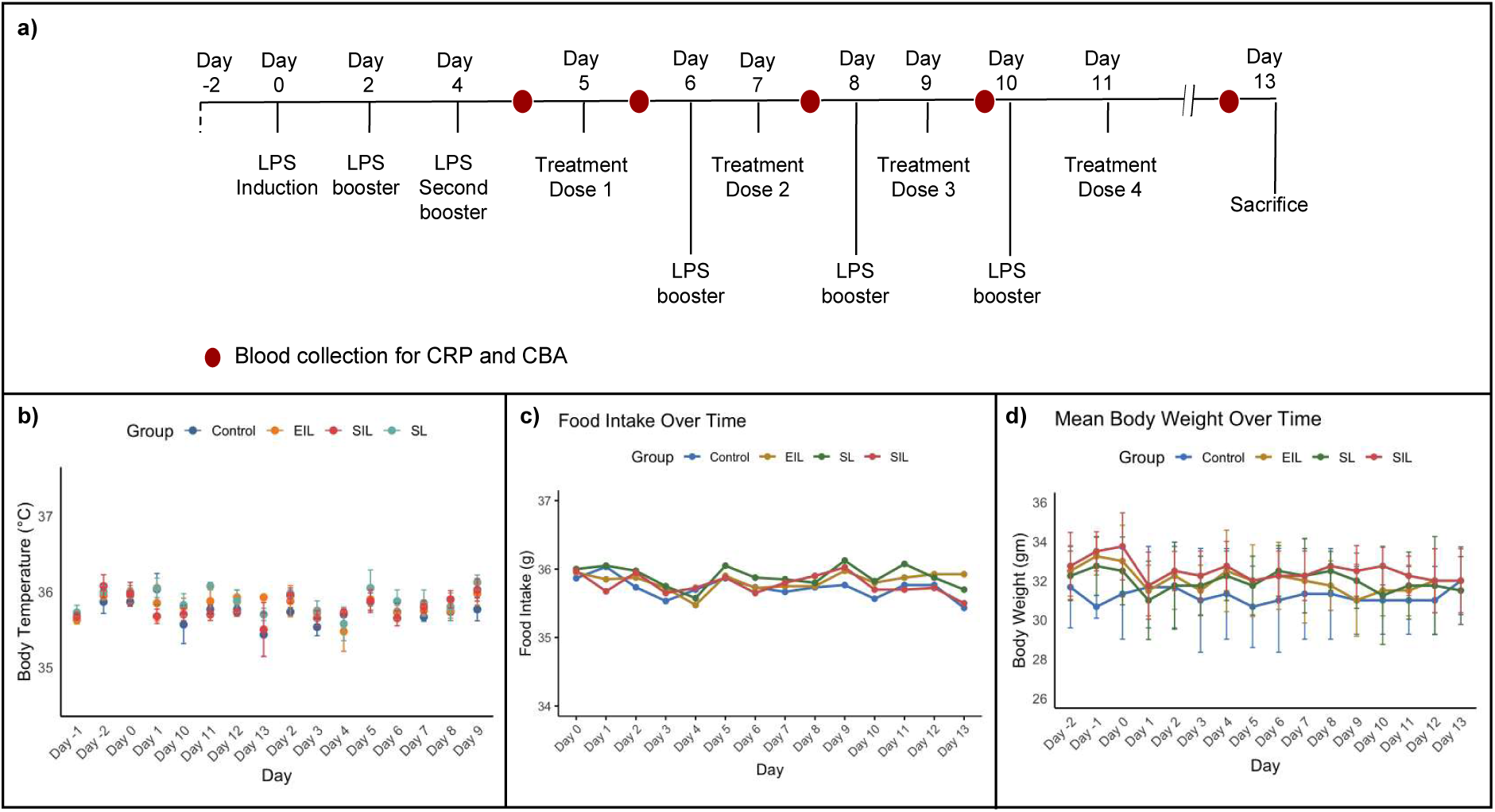
(a) Experimental workflow depicting the treatment regimen in four groups: Control, EIL (empty immunoliposome), SL (siRNA-liposome), and SIL (siRNA-immunoliposome), (b) Body temperature measurements across 14 days showed mild day-to-day fluctuations but remained within the physiological range in all groups, (c) Food intake over time displayed transient variations, with no consistent trend of reduction or increase attributable to treatment, (d) Body weight trajectories demonstrated minor fluctuations across days, with overlapping error margins between groups, suggesting that repeated dosing did not produce overt systemic toxicity. Data are presented as mean ± SD (n = 4 mice per group).

### 2.10 Systemic Inflammatory and Immunological Profiling of SIL Treatments

Systemic inflammatory responses were first assessed by quantifying CRP levels (Figure 8a). In the induction group, CRP levels increased nearly 10-fold over control at dose 1 (≈180 µg/mL vs ≈18 µg/mL, ****p < 0.0001). This elevation persisted across subsequent boosters, reaching ≈200–220 µg/mL (dose 2–3), representing a sustained 10–12-fold increase relative to baseline. The EIL group mirrored this profile, confirming that the liposomal vehicle had no intrinsic anti-inflammatory effect. In contrast, SL treatment reduced CRP to ≈90 µg/mL (≈5-fold above control) at dose 1 and progressively suppressed CRP by doses 3–4, where levels fell to ≈40– 50 µg/mL. SIL treatment exhibited the strongest effect, with CRP restricted to ≈25–35 µg/mL across all doses, corresponding to only a 1.5–2-fold elevation over control and a ∼6–8-fold reduction compared to induction.

**Figure 8.**
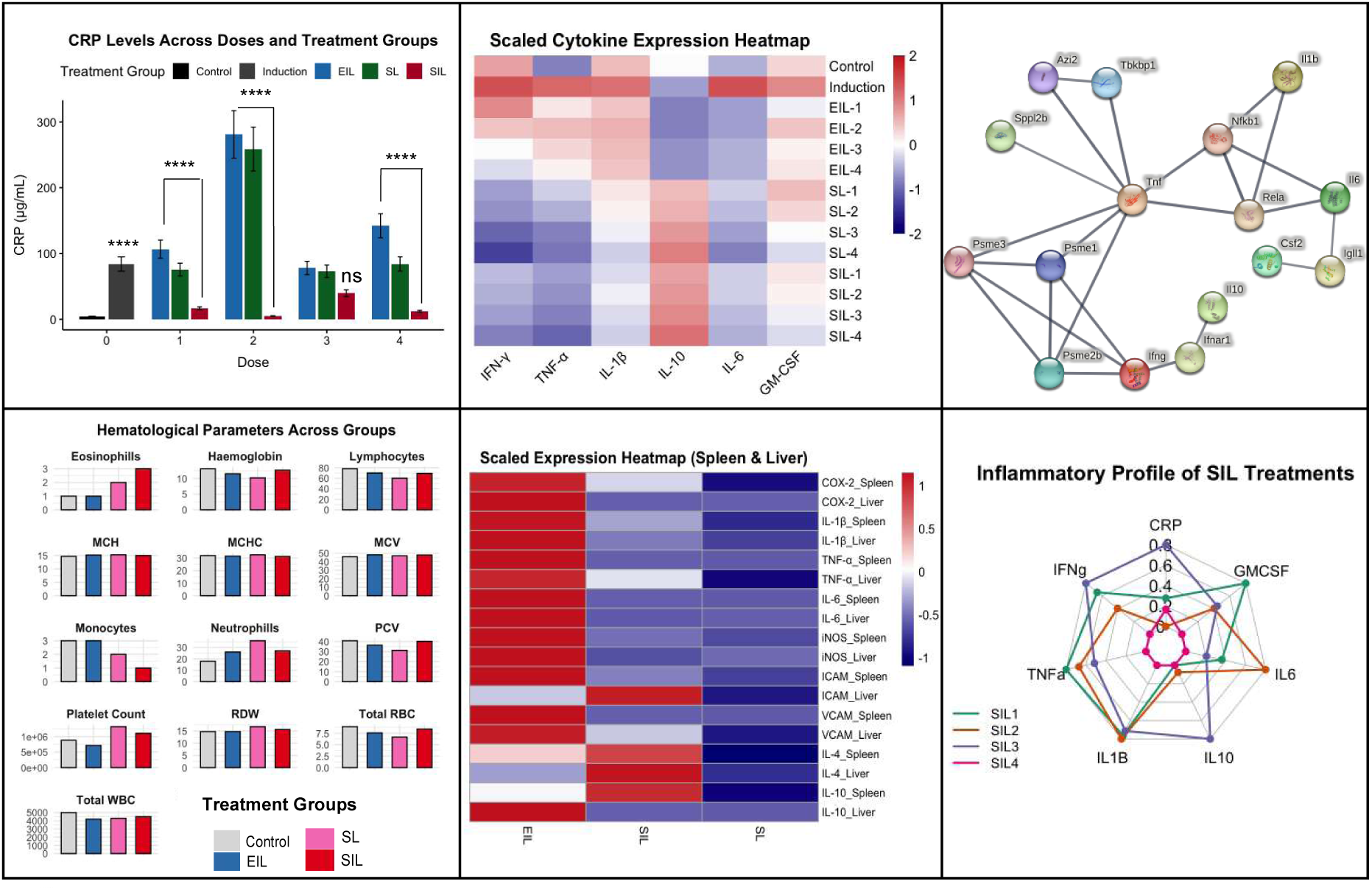
Evaluation of systemic inflammatory and immunological responses following siRNA-liposome (SL) and siRNA-immunoliposome (SIL) treatments. (a) C-reactive protein (CRP) levels across groups and doses. The induction group showed significant CRP elevation compared with the control, while EIL mirrored the induction profile. In contrast, SL and SIL treatments resulted in markedly attenuated CRP responses across booster doses, with SIL maintaining the lowest CRP levels. Data are mean ± SD; ****p < 0.0001, ns: not significant, (b) Heatmap of scaled cytokine expression (serum) revealed robust induction of IFN-γ, TNF-α, IL-1β, and IL-6 in the induction and EIL groups, while SL showed partial suppression and SIL exhibited broad downregulation of proinflammatory cytokines with relative preservation of IL-10, (c) STRING protein–protein interaction network for key inflammatory cytokines identified from expression profiling, highlighting central roles of TNF, IL-1β, and IL-6 nodes with their regulatory connectivity, (d) Hematological parameters across groups showed no significant deviation from baseline ranges, confirming the absence of systemic hematotoxicity in SL and SIL groups, (e) Heatmap of scaled expression of inflammatory mediators in spleen and liver. SIL markedly reduced COX-2, IL-1β, TNF-α, IL-6, iNOS, and adhesion molecules (ICAM, VCAM) compared with SL and EIL, with a trend toward higher IL-10 expression, suggesting suppression of systemic and tissue-level inflammatory responses, (f) Radar plot of inflammatory profile across SIL doses showed consistent downregulation of CRP, TNF-α, IL-1β, and IL-6, with relative sparing of IL-10, indicating a dose-consistent anti-inflammatory profile.

Serum cytokine heatmap analysis (Figure 8b) revealed distinct immunomodulatory signatures. Induction and EIL groups displayed robust upregulation of IFN-γ, TNF-α, IL-1β, and IL-6, each increased by approximately 2-fold relative to control. SL attenuated these cytokines to a moderate extent, reducing IL-1β and TNF-α by ≈30–40%. Notably, SIL treatment reduced proinflammatory cytokines by ≈60–70% compared to induction, while maintaining IL-10 expression at near-control levels. Thus, SIL achieved a dual effect of suppressing effector cytokines while preserving immunoregulatory signaling. STRING network analysis (Figure 8c) confirmed that the dominant inflammatory hubs (TNF, IL-1β, IL-6) were effectively suppressed under SIL, highlighting its impact on key proinflammatory axes.

Hematological analysis across groups (Figure 8d) demonstrated stable counts of leukocytes, erythrocytes, and platelets. No significant deviations were observed in eosinophils, lymphocytes, or monocytes compared with control. Erythrocyte indices (MCV, MCH, MCHC, RDW, PCV) and total WBC remained within physiological ranges across all groups. These results indicate that neither SL nor SIL imparted hematotoxicity, further supporting their systemic safety.

At the tissue level, expression profiling of spleen and liver (Figure 8e) showed that induction and EIL groups were characterized by elevated COX-2, IL-1β, TNF-α, IL-6, and iNOS, with values approximately 1.5–2-fold higher than control. SL partially reduced these mediators (∼30–40% decrease). In contrast, SIL caused near-complete suppression, lowering inflammatory gene expression by ≈60–80% across both spleen and liver. Adhesion molecules (ICAM, VCAM) followed a similar trend, indicating reduced leukocyte recruitment potential under SIL treatment. Interestingly, IL-10 expression in SIL-treated spleen and liver was ≈1.3– 1.5-fold higher than in control and ≈2-fold higher than in induction, suggesting that SIL skews responses toward an anti-inflammatory phenotype.

Radar plot analysis (Figure 8f) across SIL doses reinforced this conclusion. CRP, TNF-α, IL-1β, and IL-6 levels were consistently suppressed by >60% compared with induction across all booster doses, while IL-10 remained stable or moderately elevated. This reproducibility indicates that SIL exerts a dose-consistent anti-inflammatory effect, maintaining efficacy even under repeated administrations. Overall, SIL significantly attenuated systemic inflammation, reducing CRP by up to 8-fold and proinflammatory cytokines by 60–70%, while enhancing IL-10 by ≈2-fold. These effects were observed both in circulation and at tissue sites (spleen, liver), without altering hematological homeostasis. The combined evidence positions SIL as a potent immunomodulatory platform capable of suppressing excessive inflammatory responses while preserving regulatory pathways.

### 2.11 Histopathological Assessment of Liver and Spleen Morphology

Histological analysis further corroborated the biochemical and molecular findings, highlighting the tissue-level consequences of induction and the protective effects of SIL treatment (Figure 9). Notably, the EIL group showed marked signs of inflammation in most organs, while the SIL treatment restored the tissue architecture. In the liver H&E sections, the induction (EIL) group exhibited disrupted hepatocyte morphology with cytoplasmic vacuolation and immune cell infiltration around the portal region (red arrows). Compared with normal control, where hepatocytes retained polygonal morphology and intact sinusoids, EIL displayed severe architectural distortion, confirming ongoing inflammatory and degenerative processes. SL treatment produced partial improvement, with moderate preservation of hepatocyte structure, whereas SIL restored liver tissue architecture almost to control levels, with only minimal infiltration visible. Periodic Acid Schiff (PAS) staining of liver revealed enhanced glycogen uptake in the EIL group, accompanied by distortion in hepatocyte arrangement (yellow arrows). This indicates metabolic stress and altered carbohydrate storage. SIL treatment significantly normalized PAS uptake, resembling the control phenotype, whereas SL showed only partial restoration. Masson’s Trichrome (MT) staining, specific for staining the Type I collagen deposition and observing the levels of fibrosis under inflammatory conditions, demonstrated extensive collagen deposition in the EIL group, particularly around portal veins and extending into inter-hepatocyte spaces (green arrows). This reflects fibrotic remodelling consistent with chronic inflammatory stress. Collagen deposition in EIL appeared 2–3 fold higher than in control. SL treatment reduced collagen accumulation modestly, while SIL markedly suppressed fibrosis, with only minimal perivascular collagen observed, suggesting a strong anti-fibrotic effect.

**Figure 9.**
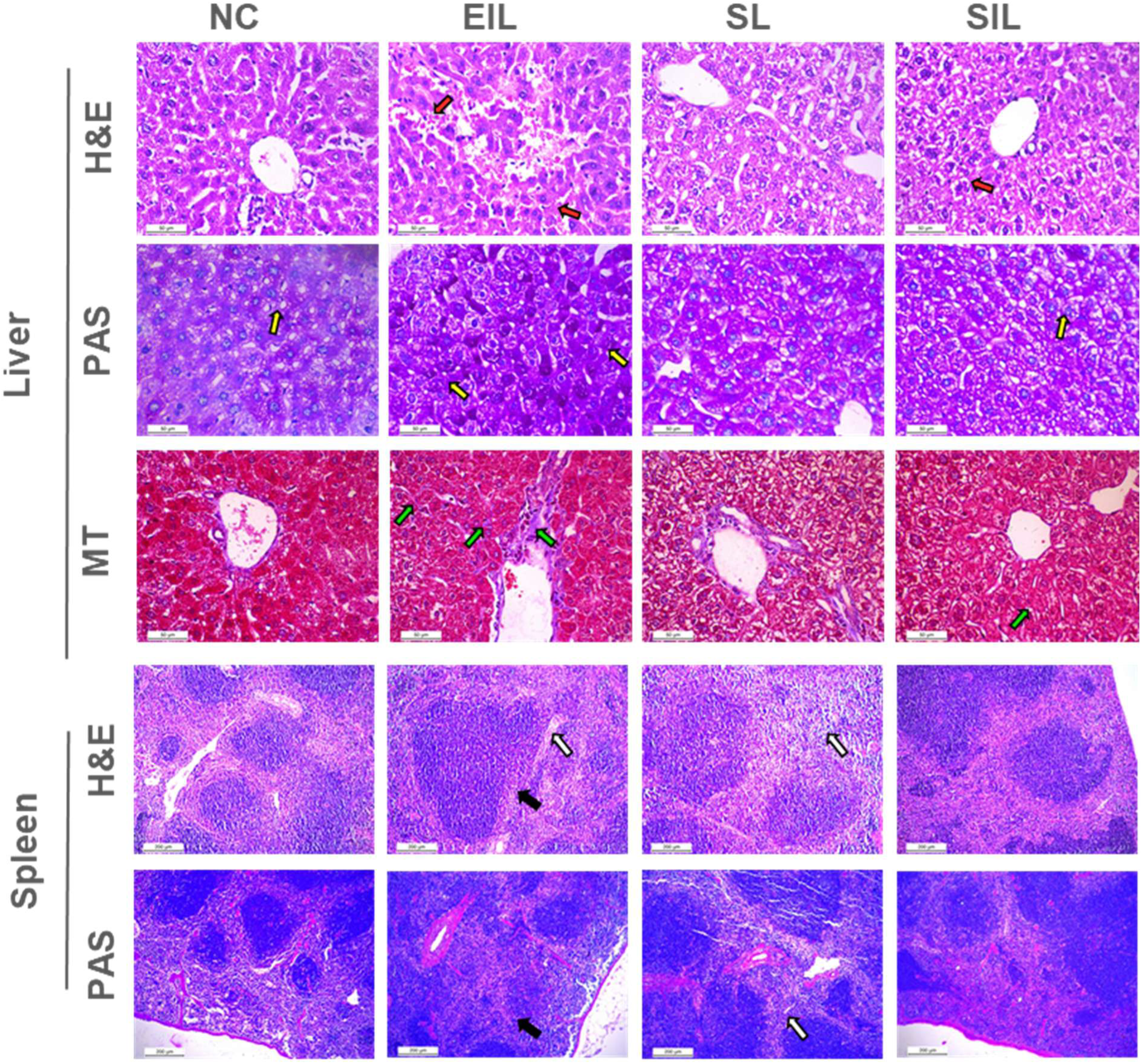
Pictorial representation of morphological changes observed at the tissue level: First panel indicates the H&E staining of Liver, with disrupted hepatocyte morphology and immune cell infiltration in EIL group (red arrows). The hepatocyte morphology is restored after SIL treatment (red arrows). PAS staining of the liver shows an enhanced uptake in the EIL group compared to control and SIL, along with distortion in hepatocytes (yellow arrows). MT staining of the liver showed high collagen deposition around the portal veins (stain uptake in purple) as well as between hepatocytes (green arrows). The SIL group shows less collagen deposition compared to the EIL group. The H&E and PAS staining of the spleen demonstrated the disappearance of the marginal zone between the red and white pulp (Black arrows) in the EIL group, along with degeneration of lymphoid follicles (White arrows). The marginal zone appears to be restored after SIL treatment. Images were acquired at 40X for the liver and 10X for the spleen.

In the spleen H&E sections, the EIL group showed loss of the marginal zone between red and white pulp (black arrows), along with degeneration of lymphoid follicles (white arrows). This structural breakdown reflects impaired splenic organization and immune dysfunction under chronic inflammatory burden. SL treatment resulted in partial recovery, but the marginal zone and follicular architecture were not completely preserved. In contrast, SIL restored both the marginal zone and follicle integrity, with overall tissue organization closely resembling that of the normal control. PAS staining of the spleen further confirmed these findings: the EIL group demonstrated disturbed parenchymal architecture with disrupted marginal zones and abnormal glycogen uptake, whereas SIL treatment largely re-established better histological features.

The histopathological changes in the kidney, particularly the increased capsular space and shrunken glomeruli, were suggestive of the abnormal changes in the Kidney’s filtration unit in the EIL group. A high level of leukocyte infiltration, tubular necrosis and disrupted cell architecture was also observed in the EIL group, suggestive of the impaired kidney functioning under chronic inflammatory conditions. Comparatively, after SIL treatment, the capsular space was restored along with minimal congestion and preserved glomeruli architecture. Similarly, the collagen and erythrocyte deposition around the glomeruli and the tubules also decreased in the SIL treated group compared to the EIL group, as shown in Figure 10 (blue arrows). The PAS staining of the kidney tissue also exhibits higher intensity in the mesengial matrix of the glomeruli in EIL group, along with disrupted basement membrane, both of which are relatively alleviated in the SIL treated group. Additionally, the brush border epithelium of the tubules in the kidney (stained magenta) also appear healthy in the SIL treated group. These observations suggest the anti-inflammatory effects of the SIL treatment on the kidney after LPS induced chronic inflammation. This anti-inflammatory effect was also observed in the H&E staining for lung tissue, where there were less infiltrates in the bronchiolar space in SIL group compared to EIL group. The infiltrates around the bronchioles are a prominent sign of lung inflammation and can lead to obstruction of the alveolar air spaces. The EIL group also exhibited high PAS uptake and disruption in the lung parenchyma, which was restored in the SIL treatment group. To summarize, the histological analysis reveals the potential of SIL treatment in restoring tissue changes associated with chronic inflammation.

**Figure 10.**
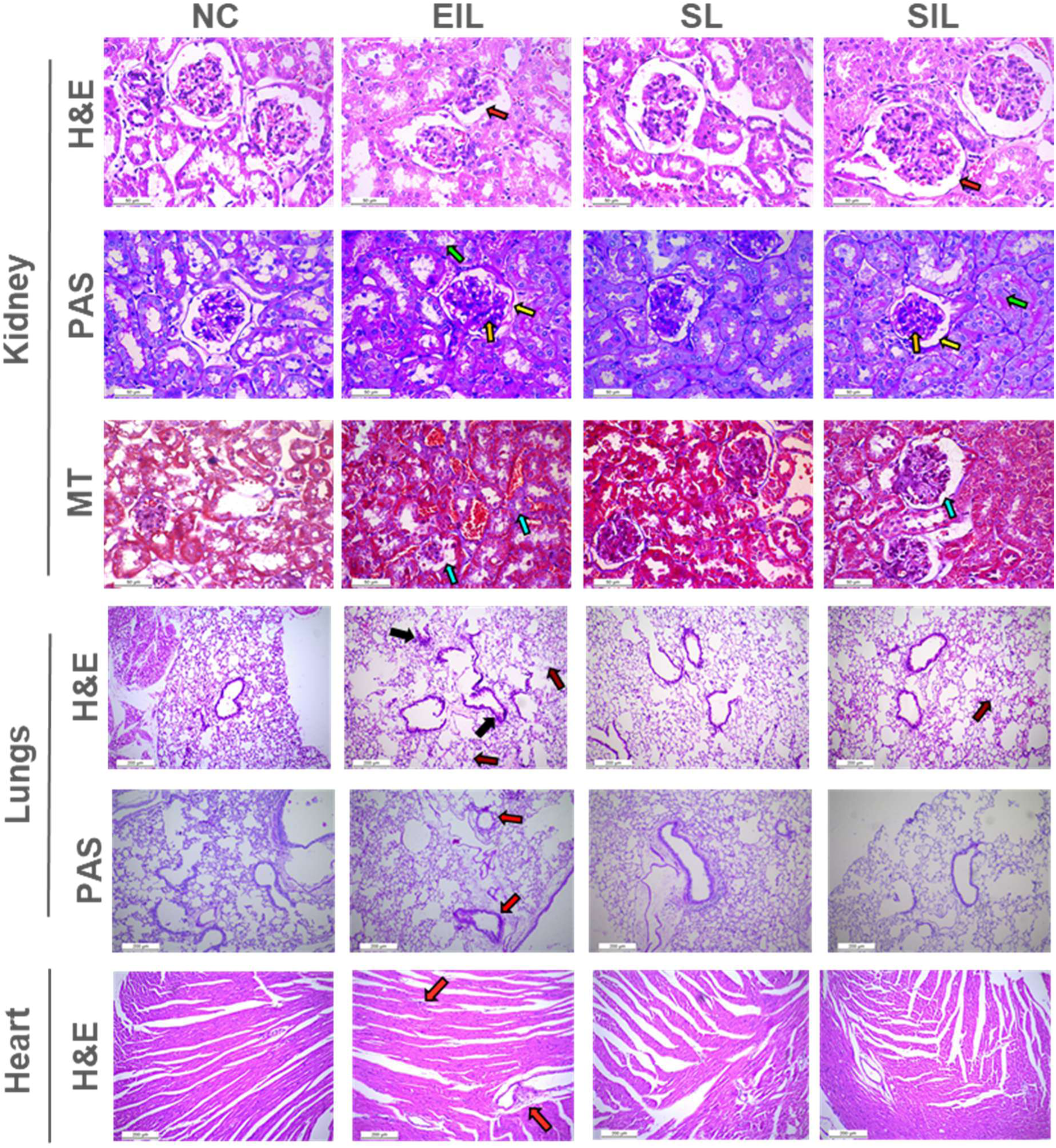
Pictorial representation of morphological changes observed at the tissue level: Disruption in the glomerular space (red arrows) was observed in the EIL group after H&E staining. Glomerulus structure appeared to be restored to normal in the SIL group (red arrows). The PAS staining of the kidney revealed high PAS uptake in the mesangial matrix in the glomeruli of the EIL group (orange arrows). Uneven uptake of the stain was observed on the basement membrane, suggestive of tissue injury (yellow arrow). Uneven Magenta staining in the tubules of EIL group indicated injury in the brush border epithelium (green arrow). The brush border epithelium appeared to be intact in SIL group (green arrow). The MT stain of the kidney showed collagen deposition at the interstitial level and the glomeruli in the EIL group. (Blue arrows). Tissue samples of the lungs showed high infiltrates around the bronchioles (black arrows) and disrupted lung parenchyma (brown arrows) in H&E staining. Comparatively the parenchyma appeared to be resorted in the SIL treated group. Strong PAS staining was also observed in the lungs of the EIL group. EIL group shows myocardial tissue degeneration, comparatively restored in the SIL group. Images acquired at 40X for Kidney and heart, and 10X for Lungs.

## 3. Materials and methods

### 3.1 Chemicals

IL-1β siRNA was procured from Thermo Fisher in lyophilized form (100µM), with the following sequence-Sense “GACAGAAUAUCAACCAACA ^30^” Anti-sense “UGUUGGUUGAUAUUCUGUC ^30^” (Catalog # NM_008361); along with scramble negative control siRNA. CD44 mAb, CD80-PE (cat: 581955), and CD86-FITC (cat: 561962) were obtained from Becton Dickinson. DSPC, Soybean Phosphatidylcholine (SPC), DSPE-PEG2000 maleimide (DSPE-PEG-Mal), and DSPE-PEG amine were procured from Avanti Polar Lipids, USA. Dulbecco’s Modified Eagle’s Medium (DMEM), Roswell Park Memorial Institute 1640 Medium (RPMI-1640), Triton X 100, 6-diamidino-2- phenylindole (DAPI), , Rhodamine 6G, Cell culture tested Cholesterol, Dimethyl sulfoxide (DMSO), antibiotic and antimycotic solution 100X liquid were procured from HiMedia, Mumbai, Maharashtra, India. Fetal Bovine Serum (FBS), FTIR Grade KBr, β-mercaptoethanol, Phalloidin-Atto 488, paraformaldehyde (PFA), and Uranyl acetate (UA) were procured from Sigma Aldrich. Phosphate buffered saline (PBS), 3-[4, 5-dimethylthiazol-2-yl]-2, 5-diphenyl-tetrazolium bromide (MTT) and Lipopolysaccharide (LPS), were procured from Thermo Fisher Scientific. MCA2305GA Rat anti Mouse CD16/CD32, GAPDH (D4C6R) Mouse mAb

### 3.2 Synthesis of siIL-1β-anti-CD44-Liposomes

Liposome synthesis was performed through thin-film hydration Method ^31^, where the lipids (DSPC: cholesterol: DSPE-PEG = 45:50:5) were mixed and dissolved in chloroform, an organic solvent. The dissolved lipids in solvent were allowed to evaporate till a thin film of lipids was formed. This was achieved by subjecting the lipids to a constant temperature of 65°C and rotation under pressure through an EV11 rotary evaporator. After the thin film was formed, the lipids were hydrated with Phosphate Buffer Saline (PBS) for 2 hours. SiIL-1β (stock=100 µм) was added during hydration for making siIL-1β-Liposomes to achieve a final concentration of 400 nM. The liposome suspension was extruded 22 times through a 100nm filter to achieve small unilamellar liposome vesicles.

For anti-CD44-Liposomes synthesis, Maleimide liposomes were first synthesized with Cholesterol: SPC: DSPE-PEG-Mal in the ratio 1.25:2.5: 0.25, by thin-film hydration method. DSPE-PEG-Mal is used instead of DSPE-PEG-Amine to facilitate conjugation with the thiol group of mAb^32^. After extrusion, the post-synthesis modification of liposomes was achieved by incubating them overnight with thiolated fractions (Fractions 3, 4 and 5) of CD44 MAb, to facilitate conventional coupling of mAb to the liposome surface. All liposome formulations were stored at 4 °C.

### 3.3 anti-CD44 mAb Thiolation

Thiolation of mAb was achieved by mixing Traut’s Reagent (8 μg mL^-1^) and CD44 mAb (4 μg mL^-1^) in the ratio 1:40 mol mol^-1^, at 37 °C under magnetic stirring at 300 rpm for 3 minutes, followed by incubation at 37 °C for one hour. PD-10 desalting column was used to eliminate unreacted Traut’s reagent after thiolation; and after the first elution, PBS/EDTA buffer (5 mM, pH 8.0) was used to collect mAb fractions. Pierce™ BCA protein assay kit was used to estimate the concentration of CD44 in eluted fractions using the manufacturer’s protocol.

### 3.4 Characterization of Liposomes

#### 3.4.1 Zetasizer

The Zetasizer (Malvern Instruments Ltd. Malvern, UK) was used to measure the hydrodynamic size, Polydispersity index and ζ-Potential of the liposome formulations. Through dynamic light scattering (DLS) the size distribution of liposomes was evaluated, and their average hydrodynamic size was obtained. The sample was prepared freshly for every read by diluting 10 µL of liposome sample in 1 mL PBS (1:100 V/V). The stability in Size and ζ - Potential of liposomes was measured for 30 days, when stored at 4 °C.

#### 3.4.2 Transmission electron microscopy (TEM)

Transmission Electron Microscopy (TEM) was used to observe the morphology of liposomes at 120/100kV voltage. The sample was prepared by diluting the liposome formulation 50 times, and staining with UA. A drop of diluted liposomes was added on the carbon coated grid and stained with 1% UA. After overnight incubation, the grid was observed in TEM (JEOLJEM1400).

#### 3.4.3 Fourier Transform Infrared Spectroscopy (FTIR)

FTIR was used for examining the integrity of functional groups of liposomes. The samples were prepared by diluting in 1X PBS 10 times V/V followed by lyophilisation with KBr. They were scanned in FTIR (PerkinElmer) at a wavelength range of 4000 to 400 cm^-1^, with 11 accumulation cycles. The results were normalised and analysed using PerkinElmer Spectrum.

### 3.5 Determination of siRNA Entrapment Efficiency and Release Kinetics

The loading efficiency of liposomes and the optimum ratio of siRNA was evaluated by running a gel retardation assay of liposome formulations with different N/P ratios. The siRNA concentration in all liposomes was kept constant, and the lipids were varied in each formulation. The gel retardation assay of the liposomes was conducted for formulations before and after dialysis (to compare the effect of unbound siRNA and excess lipids left in liposome suspension). After the N/P ratio for the liposome formulations was established, the siIL-1β-Liposome release kinetic profile was performed. SiIL-1β-Liposomes were loaded in micro centrifuge tubes sealed with activated dialysis membrane, and the prepared tubes were inverted and placed overnight in a beaker containing fresh PBS (pH 6.5), under constant stirring. The ratio of liposomes to PBS was maintained at 1:50 V/V. After twelve hours, the PBS solution was collected, and the entrapment efficiency was calculated from it using the formula –

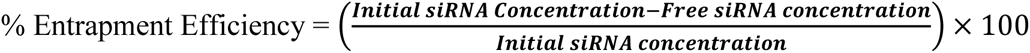

Subsequently, fresh PBS was added to the beaker and the setup was maintained for 48 hours. Hourly samples of PBS were collected from the beaker to determine the release profile of siRNA from the liposomes (1-8hr, 12hr, 24hr, and 48hr). The aliquots of the hourly samples and entrapment efficiency were analysed using the Take-3 Nucleic Acid Quantification, BioTek Synergy HT spectrophotometer. The standard curve of siRNA at different concentrations was used to calculate the released siRNA

### 3.6 Cell Culture Conditions

The macrophage cell line RAW 264.7 and T lymphocyte cell line Jurkat E6.1 were procured from National Centre for Cell Science (NCCS), Pune, India. The macrophages were maintained at 37°C in a High Glucose DMEM culture medium, and passage numbers 30 to 37 were maintained for all experiments. The T cells were maintained at 37°C in RPMI-1640 Medium and passage numbers 42-49 were used for experiments. Both cell lines were supplemented with 10% FBS and 1% Antibiotic Antimycotic Solution; and the cells were cultured in a 5% CO2 atmosphere under controlled environment conditions

### 3.7 Cell Proliferation Assay

The MTT Assay was carried out on macrophages and T cells for dosage estimation and the comparative analysis of the cytotoxicity of formulations on the cells. For macrophages, approximately 7×10^5^ cells per well were seeded in 96 culture well plates and incubated overnight at 37 °C. After successful adherence of cells on the well plate, the macrophages were stimulated with LPS (100 ng mL^-1^) for 18-24 hours. After the Macrophages were activated and achieved a pro-inflammatory Phenotype, the different liposomal formulations were added to the cells. (Bare siIL-1β and siIL-1β-anti-CD44-Liposomes). After 24 and 48 hours, the media was replaced with serum-deprived media and MTT (5 mg mL^-1^), followed by incubation for 3-4 hours. After incubation, the media was replaced with DMSO (100 μL), and the cell proliferation at 24 hours and 48 hours was quantified using the BioTek multiple plate spectrophotometer after shaking for 5 minutes and measuring the absorbance at 570 nm. For T cells, treatments with different concentrations of bare siIL-1β and siIL-1β-anti-CD44-Liposomes given to the cells and the tubes were incubated at 37°C for 24 and 48 Hours. Following incubation, the cells were given a PBS wash, re-suspended in PBS, and dispensed equally into 96 well plates and MTT (5 mg mL^-1^, 20 μL) was added to each well, followed by incubation for 3-4 hours. After incubation, formazan crystals were dissolved with DMSO (100 μL) in each well and the absorbance was measured at 570 nm using the multiplate spectrophotometer. Complete media and PBS were used as negative controls, while Triton-X- 100 was used as a Positive control.

### 3.8 Internalisation of Rhodamine Liposomes

Rhodamine (0.1 mg mL^-1^) was used to prepare Rhodamine-loaded liposomes and Rhodamine loaded anti-CD44-Liposomes for studying their cellular uptake. 50×10^3^ M0 Macrophages were seeded and activated with LPS (100 ng mL^-1^) for 18–24 h. After activation, the cells were given a PBS wash and treated with Rhodamine Liposomes and Rhodamine-loaded anti-CD44- Liposomes and allowed to incubate for 3 hours. Following incubation, the cells were fixed with PFA (4%) and stained with DAPI (0.1 mg mL^-1^) and Phalloidin (0.1 mg mL^-1^). The cells were mounted with Moviol for permanent slide preparation and observed using confocal microscope (Nikon Eclipse Ti2). A suitable objective lens with 100X magnification (with digital magnification) was used for imaging at the excitation/emission wavelengths of 350/465 nm, 495/519 nm and 546/567 nm for DAPI, FITC and Rhodamine respectively.

### 3.9 Scanning Electron Microscopy (SEM)

Scanning electron microscopy was performed to analyse the morphological changes in RAW264.7 cells after LPS activation, confirming their transition to a pro-inflammatory M1 like phenotype, and the effect of siIL-1β-anti-CD44-Liposomes on the LPS activated macrophages.

Briefly, 50×10^3^ RAW264.7 cells were seeded in 6 well plates and treated with LPS (100 ng mL^-1^), and after 24 Hours, they were treated with siIL-1β-anti-CD44-Liposomes. Later they were fixed with PFA (4%) or glutaraldehyde (2.5 %). After fixation, the cells were dehydrated by giving washes with increasing concentrations of Ethanol (25%, 40%, 60%, 80%, and 100%). After air drying and coating with gold-platinum sputtering, the cells were imaged with the Scanning Electron Microscope (JEOL JSM-6010LA)

### 3.10 Gene Expression Studies

#### 3.10.1 RNA Isolation and cDNA synthesis

Total RNA was extracted from cells and snap frozen tissue samples (liver and spleen) using the Favourogen RNA extraction kit, using the manufacturer’s protocol. RNA was isolated from untreated macrophages and T cells, LPS activated macrophages and T cells, and cells treated with 150 pм siIL-1β-anti-CD44-Liposomes for 24 and 48 hours. The isolated RNA was checked for its concentration and purity using the SYNERGY-HT multiwell plate reader (Synergy HT, Bio-Tek, USA). The RNA was used for cDNA synthesis using the Biorad cDNA synthesis kit according to the manufacturer’s protocol.

#### 3.10.2 qRT-PCR for gene expression studies

The SYBR Green Master Mix (Applied Bio systems) was used for real-time PCR of the cDNA, along with the sense and anti-sense primers specific for pathways in Macrophages and T cells. GAPDH and B-actin was used as a housekeeping gene and the gene expression was evaluated with the inactivated cells as the control. Eppendorf thermo cycler was used for performing the PCR and Quant Studio 5 was used for the quantification of the PCR Results. The PCR was performed for 40 cycles, and the quantification was done by the ΔΔCT method

#### 3.10.3 Co-Culture of Macrophages and T cells

The effect of LPS activated Macrophages on T cell co-stimulation was studied by the co-culture of Macrophages and T cells. Briefly, Macrophages were activated by treatment with LPS, and after 24 hours a PBS wash was given to remove excess LPS. Since activated macrophages release certain cytokines and exhibit elevated co-stimulatory markers, they have the potential to cause changes in T Cell Programming. This was studied by co-culturing T cells with macrophages. The changes in gene expression of T cells under the influence of activated macrophages and after treatment with siIL-1β-anti-CD44-Liposomes were studied through qPCR. GAPDH was used as the housekeeping gene and the cDNA from T cells co-cultured with inactivated macrophages was used as a control. The RNA Isolation, cDNA synthesis, and RT-PCR were performed with the procedure described in Section 3.10.1 and 3.10.2.

### 3.11 Western Blotting

RAW264.7 were seeded at a density of 5×10^4^ cells per well, and after 24 and 48 hour treatment with 150 pм siIL-1β-anti-CD44-Liposomes, the cells were lysed for protein extraction using RIPA lysis buffer. Protein concentration was estimated using the BCA protein estimation kit, and SDS-PAGE and western blot of the samples was carried out. 10% resolving gel and 4% stacking gel was prepared for SDS-PAGE of the samples, and 80 µg/µL sample was loaded per well. 10–175 kDa protein ladder (PageRular, Thermo Scientific) was used as a molecular weight marker. GAPDH (catalog # D4C6R) primary mAb was used as the housekeeping control. IL-1-beta (catalog # 3A6) Mouse mAb, NLRP3 (catalog # D4D8T) Rabbit mAb, and TNF-alpha (catalog # D2D4) XP(R) Rabbit mAb (Mouse Specific) primary antibodies were procured from Cell Signalling Technology. Chicken anti-mouse IgG H&L (HRP) (Abcam, Cambridge, UK) and chicken anti-rabbit IgG H&L (HRP) (Abcam, Cambridge, UK) were used as the secondary antibodies. The blot was developed using a Bradford Clarity Max^TM^ Western ECL substrate and observed in the ImageQuant LAS500 gel documentation system. The statistical analysis was carried out using ImageJ 1.52a software. Two-way ANOVA was carried out to calculate the significance using GraphPad Prism (version 5; La Jolla, CA, USA) software. The following adjustments were maintained across all blots – Brightness: 13 and Contrast: 32

### 3.12 Cell surface Markers profiling

RAW264.7 were seeded at a density of 1-5×10^6^ cells per well and different concentrations of siIL-1β-anti-CD44-Liposomes treatments were given after 18-24 hours of LPS activation. Following 24 hours of liposome treatment, the cells were first incubated with F_c_ Block and then with CD80/CD86 cocktail (0.1-1µg/mL). The cells were harvested after 30 minutes of mAb incubation and acquired in Beckman Coulter Cytoflex Flow Cytometer.

### 3.13 Animal procurement and maintenance

Healthy C57 BL/6 mice were procured from Zydus Research Centre, Ahmedabad, India, and the experiments were performed at LM College of Pharmacy (LMCP), Ahmedabad. The mice were kept in standard conditions under the CPCSEA in an animal house of LMCP, with a relative humidity of 60 ± 5% and a temperature of 25 ±2 °C. A 12h:12h light-dark cycle was also maintained for these animals. Balanced rodent food pellets and water were provided. All experiments were reviewed and accepted by the Institutional Animal Ethics Committee. (LMCP/IAEC/24/0044)

### 3.14 Dosage and Induction

The animals were allowed to acclimatise for 7 days before LPS induction. LPS (1.5 mg/kg BM) was given on alternate days to induce inflammation through Intraperitoneal injection, based on Ramirez et al’s model of systemic inflammation ^33^. After three LPS doses, treatment was administered via Intraperitoneal injection. The treatment regimen is provided in Figure 7 (a) of the manuscript. The body weight, food intake, and temperature (using a Perfecxa infrared thermometer) were measured throughout the duration of the study. After two weeks, the animals were sacrificed, and the Spleen, kidney, liver, heart, and lungs were isolated for further analysis. The organs were fixed in 10% neutral buffered formalin for sectioning and histological examination. The spleen and liver were snap frozen for RNA isolation and gene expression studies. Blood CBC profiling was done immediately after sacrifice at Sukoon Pathology Laboratory, Ahmedabad.

### 3.15 Blood collection, Serum Isolation, and CRP measurement

Retro-orbital plexus blood collection under di-ethyl ether anaesthesia was performed after 24 hours of treatment to measure the levels of C - reactive protein in the treatment groups. 30 minutes after blood collection, the serum was isolated from the blood by centrifuging at 1500xg for 25 minutes. The CRP was measured using a Turbilatex commercial kit using the manufacturer’s protocol (Beacon Diagnostics Ltd, India).

### 3.16 CBA analysis

The isolated serum samples were analysed for levels of secretory cytokines using LEGENDplex Mouse Inflammation Panel, by the manufacturer’s protocol for V-bottom plates and serum samples. FL10-A (B610) was used as a reporter channel for PE, and FL-17 (R-660) was used as a classifying channel for APC. Channel A measured cytokines IFN-*γ* Capture bead A6 (Cat - 740153) and TNF-ɑ Capture bead A7 (Cat - 740154). Channel B measured IL-1**β** Capture bead B2 (Cat - 740157), IL-10 Capture bead B3 ( Cat - 740158), IL-6 Capture bead B4 (Cat - 740159), and GM-CSF Capture bead B9 (Cat - 740163). The standard curve and concentration for the calibrator beads are given in the Supplementary information.

### 3.17 Staining and visualization

The fixed tissue samples were sectioned and stained with haematoxylin and eosin for histopathological analysis. The sectioning and staining were performed at Sukoon Pathology laboratory, Ahmedabad. PAS and MT staining was performed at Unipath Laboratory, Ahmedabad. The visualization was performed through the DM2500 Leica microscope, with constant brightness and exposure across panels.

### 3.18 Statistics and reproducibility

The statistical analysis was performed through GraphPad Prism (version 5; La Jolla, CA, USA).The experiments were performed in biological triplicates. The statistical analysis was done using GraphPad Prism analysis tool through Kruskal–Wallis test, t-test and two-way ANOVA and represented as mean ± SD. The P-values and their significance were as follows. * p < 0.05, ** p < 0.01, *** p < 0.001, **** p < 0.0001, respectively.

## 4. Conclusion

In conclusion, this study tries to unravel the intricate dynamics associated with cytokine mediated chronic inflammation, and how CD44-mediated delivery of siIL-1β can work as an effective therapeutic against the same. SiIL-1β-anti-CD44-Liposomes (SIL) were successfully formulated through thin-film hydration method, with well-preserved size and stability over a duration of 10 months, whenever randomly measured. The liposomes also exhibited exceptional entrapment efficiency, followed by a sustained release of the siRNA in 24 hours. Targeted delivery of liposomes through CD44 is observed in LPS-activated Macrophages, underscoring the potential therapeutic implications for precision therapy in conditions associated with macrophage mediated inflammation. The studied molecular mechanisms also display the anti-inflammatory and immune regulation effects of siIL-1β-anti-CD44-Liposomes on *in-vitro* systems through multifaceted cascade of events. The anti-inflammatory properties observed in Macrophages after treatment underscored the siIL-1β-anti-CD44-Liposome’s ability to regulate the macrophage mediated inflammation at a minimal dosage of 150 pм. Additionally, their effects on the macrophage-T cells crosstalk reveal their nuanced effects on T cell effector functions associated with Antigen presentation, suggesting their potential for precise therapeutic interventions in conditions characterized by heightened inflammation. Moreover, the in-vivo studies revealed the promising potential of the SIL treatment in alleviating chronic inflammation, as observed through the significant changes in the levels of pro-inflammatory cytokines and CRP. The ability of the SIL on the tissue level also promises them as a better therapeutic agent for chronic inflammatory conditions.

## Author contributions

HS: Experimentation, Experiment design, Data analysis and Manuscript writing; SN: Conceptualization, Experiment design, Manuscript writing, Data analysis; MP: Figure preparation and review of manuscript; DB: Manuscript review and data analysis AK: Funding acquisition, conceptualization, Experiment design and Manuscript review.

## Acknowledgements

The authors sincerely thank Gujarat State Biotechnology Mission (GSBTM) (URBASASE22C5/GSBT/23-24_AK_12.26) for providing financial support. HS would like to acknowledge Ahmedabad University for providing the fellowship to assist her research.

## Data availability statement

Data will be made available on request.

