## Supplementary file for "Preclinical evaluation of Targeted IL-1β Knockdown via CD44-Immunoliposomes: A Nano-therapy against the Inflammatory Microenvironment"

*Corresponding Author:

Ashutosh Kumar,

Associate Professor

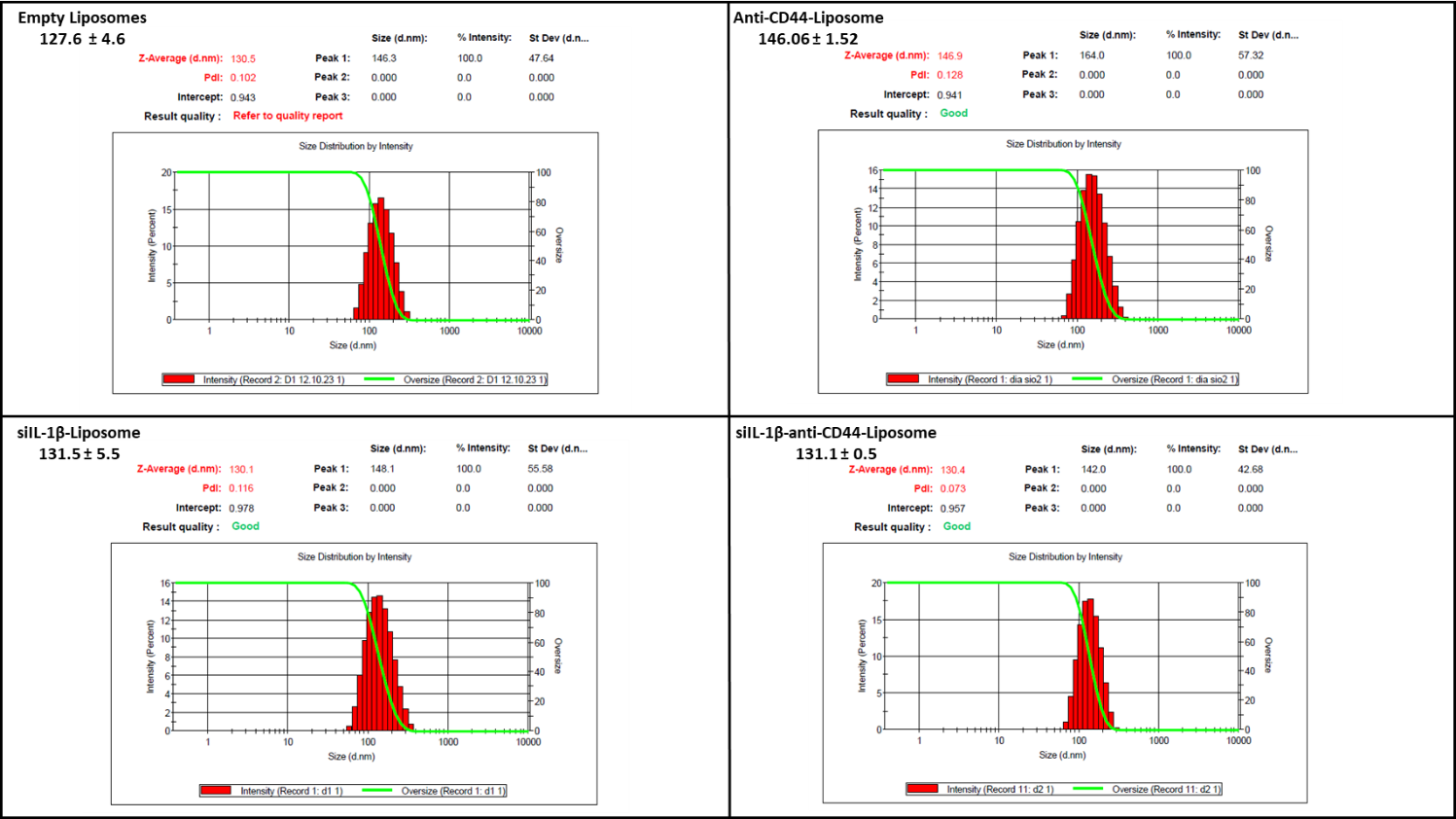

*Figure S1: Size distribution by intensity of different liposome formulations.*

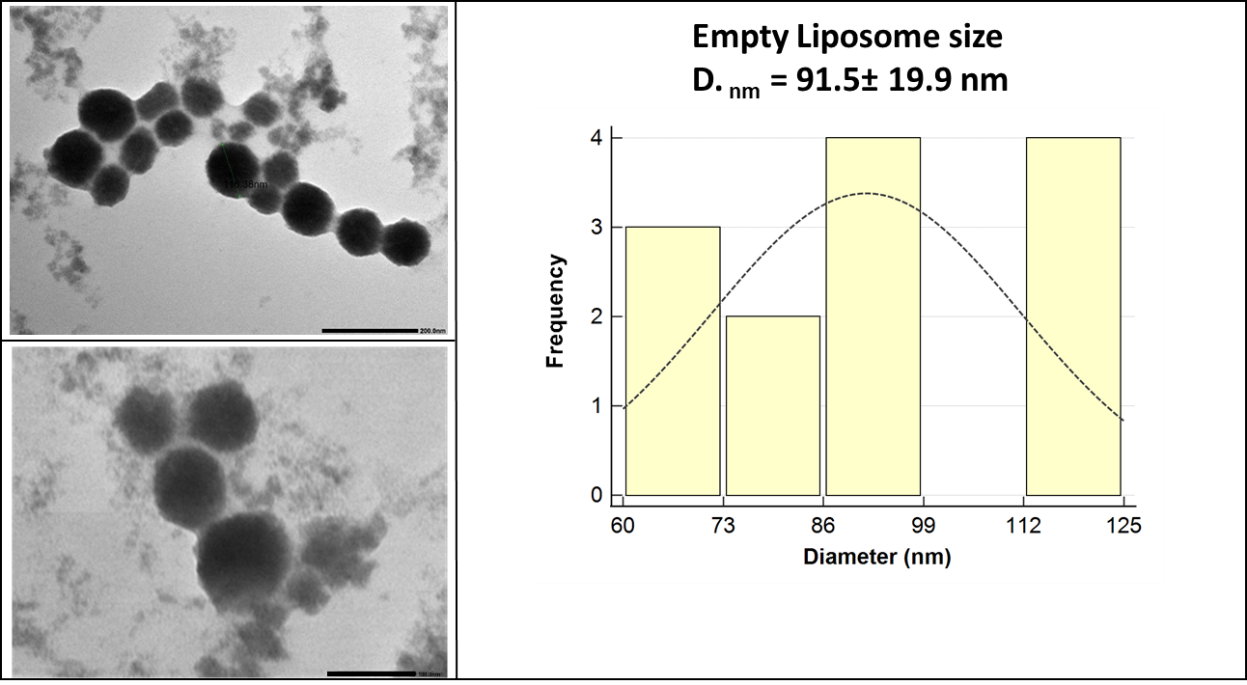

*Figure S2: TEM images and size-distribution histogram of liposome formulation before mAb conjugation.*

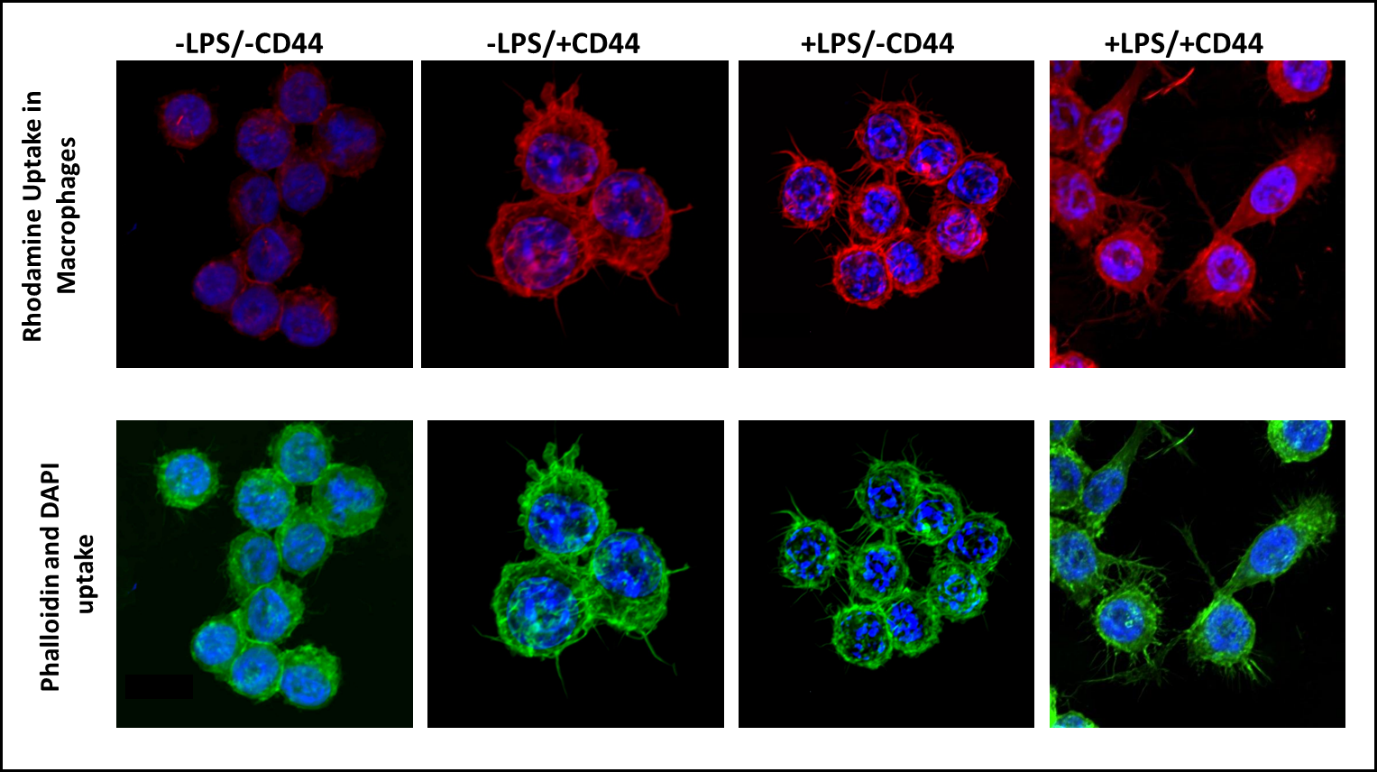

*Figure S3: Pictorial representation of enhanced uptake of Rhodamine after CD44 mAb functionalization on liposomes as observed through Confocal Microscopy at 100X (digital magnification). –LPS/+LPS represent the M0 and M1 phenotype of cells. –CD44/+CD44 represents the liposomes untagged with CD44 mAb versus tagged with CD44 mAb, respectively; Phalloidin and DAPI are used at counter stains.*

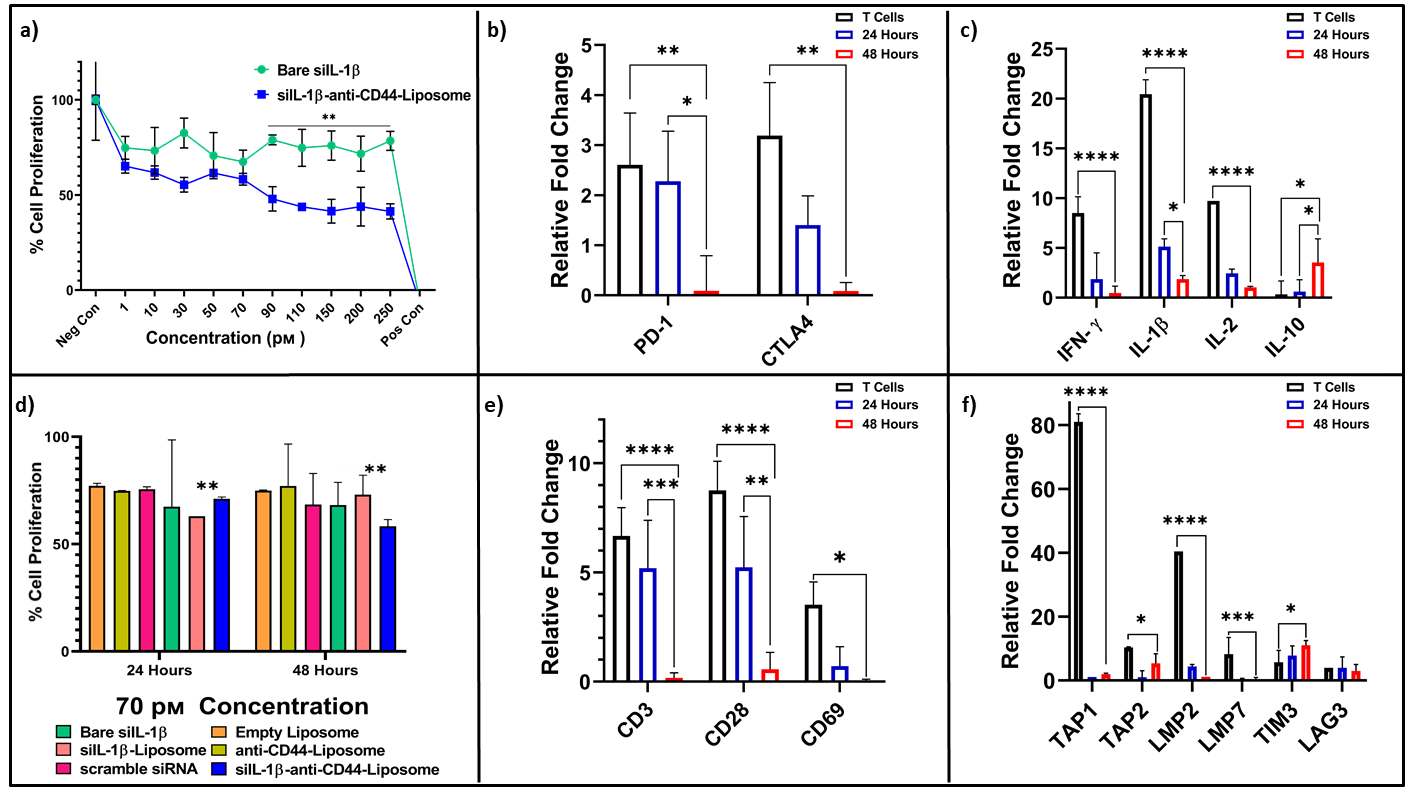

*Figure S4: Graphical representation of the change in cell proliferation of Jurkat E6.1 Cells. a) Represents the cytotoxicity induced by siIL-1β-anti-CD44-Liposomes as compared to bare-siIL1β at 48 hours duration d) Represents the negligible cytotoxicity of various formulations at 70pᴍ concentration, the safe range determined through IC_50_ Calculation b-c) and e-f) Analysis of gene expression on Jurkat E6.1 cells after 24 and 48 hours of siIL-1β-anti-CD44-Liposomes treatment. The p-values for these graphs are represented as * for p < 0.05, ** for p < 0.01, and *** for p < 0.001 using two-tailed t-test. All experiments were performed in triplicates.*

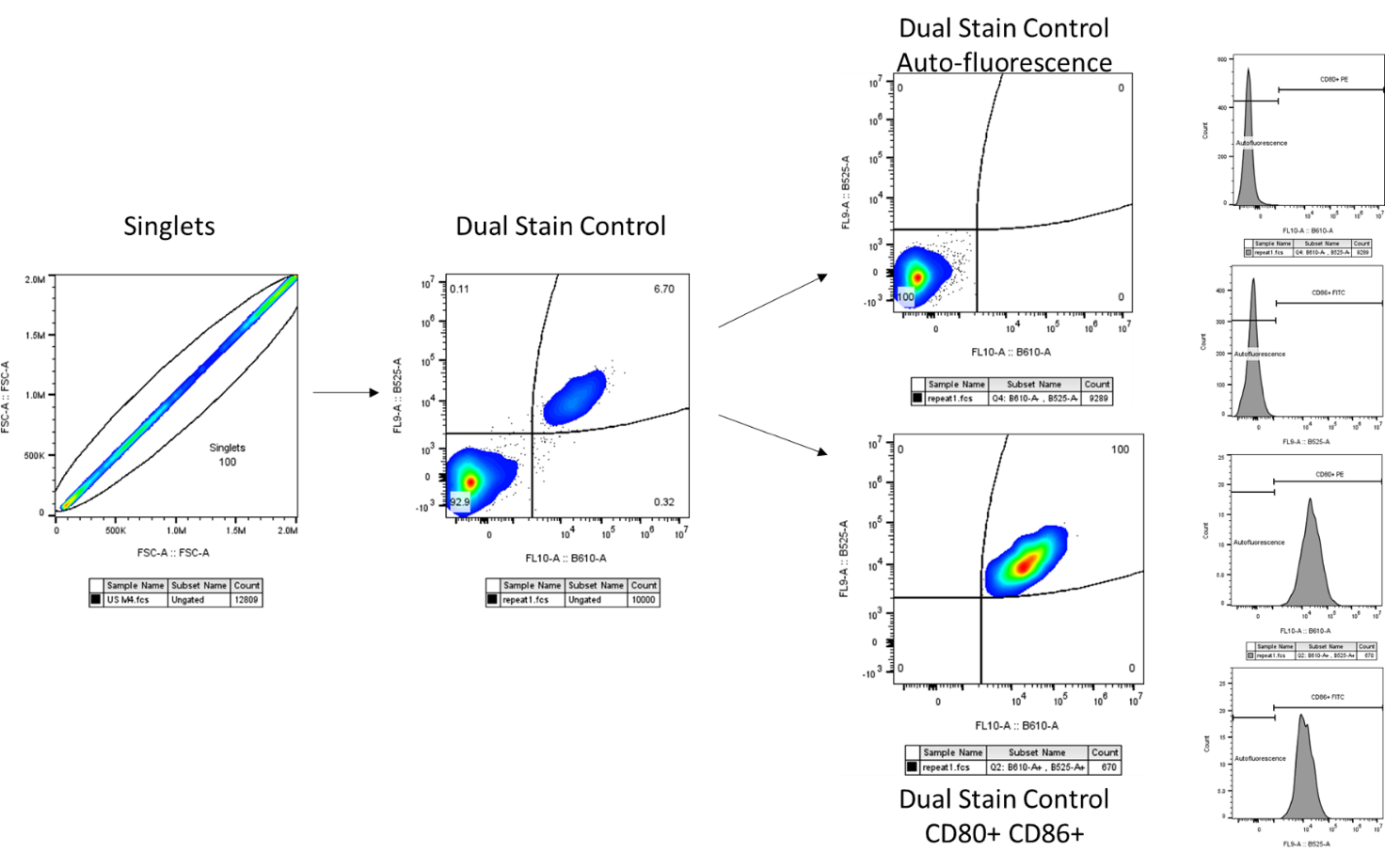

*Figure S5: The gating strategy applied for Cell surface markers analysis, with dual stained cells as control.*

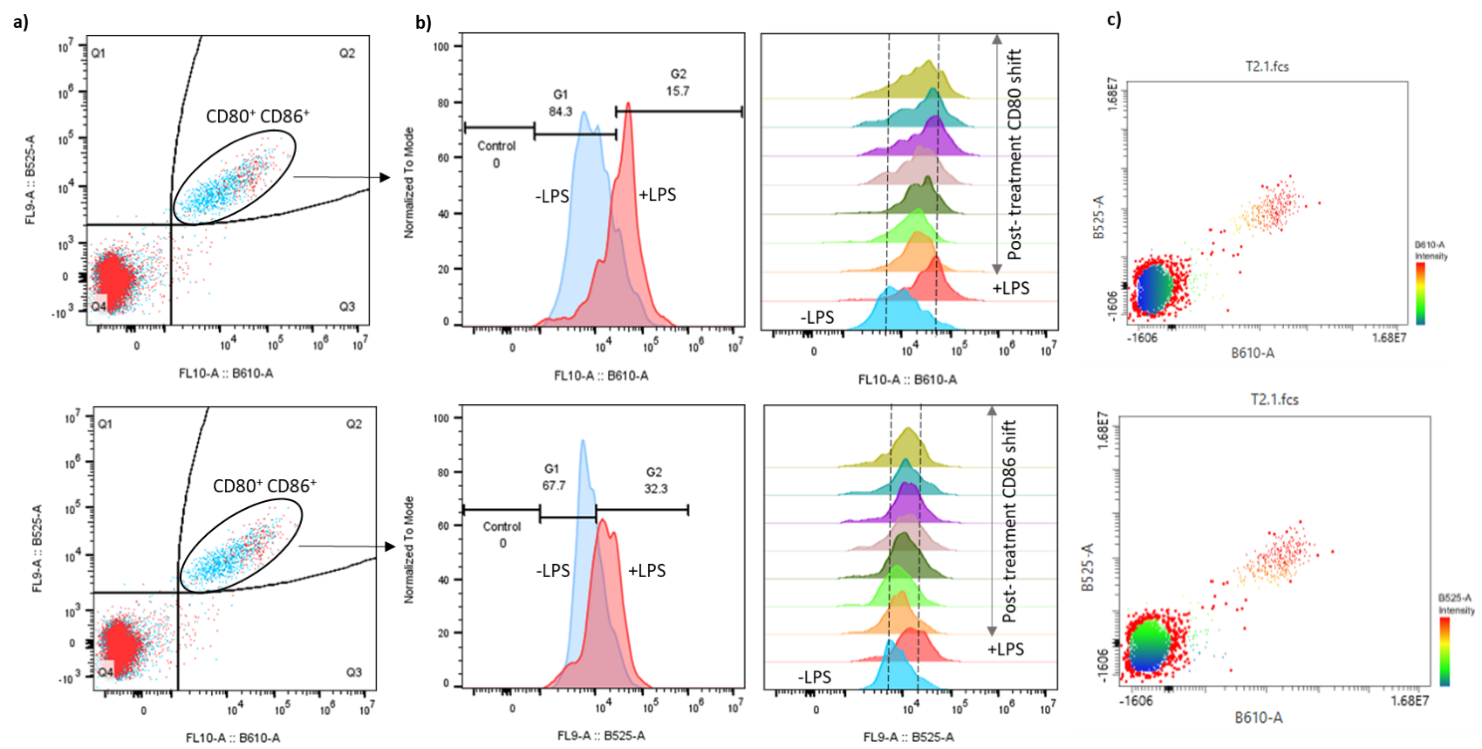

*Figure S6: The gating strategy applied for cell surface markers analysis.a) represents the dual stained population b) represents the shift in the CD80/86 levels in different treatments c) represents the dot plot of CD80/86 population at treatment concentration 50 pᴍ. The heat map depicts the intensity of CD80/86 expression.*

| ***Gene*** | ***Forward primer (5′−3′)*** | ***Reverse primer (5′−3′)*** |
| --- | --- | --- |
| *IL1β (Mouse)* | *TGGTGTGTGACGTTCCCATT* | *CAGCACGAGGCTTTTTGTTG* |
| *COX-2* | *TCCAAATGAGATTGTGGGAAAATTGCT* | *AGATCATCTCTGCCTGAGTATCTT* |
| *IL-6* | *CCGGAGAGGAGACTTCACAG* | *GAGCATTGGAGGTTGGGGTA* |
| *IL-4* | *ACCTTGCTGTCACCCTGTTC* | *CAGTGTTGTGAGCGTGGACT* |
| *PD-1* | *AAGTCAATGCCCCATACCGC* | *CTCTTCCCACTCACGGGTTG* |
| *iNOS* | *TCCTGGACATTACGACCCCT* | *CTCTGAGGGCTGACACAAGG* |
| *TNF-α* | *AGCCCACGTCGTAGCAAACCAC* | *AGGTACAACCCATCGGCTGGCA* |
| *IL-10* | *GAGTGAAGACCAGCAAAGGC* | *CAAGGAGTTGCTCCCGTTAG* |
| *ICAM-1* | *AGGTATCCATCCATCCCACA* | *AGTGTCTCATTCCCACGGAG* |
| *VCAM-1* | *CGGTCATGGTCAAGTGTTTG* | *GAGATCCAGGGGAGATGTCA* |
| *IL1β (Human)* |  |  |
| *PD-1* | *CGTGGCCTATCCACTCCTCA* | *ATCCCTTGTCCCAGCCACTC* |
| *CTLA4* | *ACGGGACTCTACATCTGCAAGG* | *GGAGGAAGTCAGAATCTGGGCA* |
| *CD3 (T Cell Co-Receptor):* | *CATTGCTTTGATTCTGGGAACTGAATAGGAGGA* | *GGCTGCTCCACGCTTTTGCCGGAGACAGAG* |
| *CD28 (Co-stimulatory Molecule):* | *GAGAAGAGCAATGGAACCATTATC* | *TAGCAAGCCAGGACTCCACCAA* |
| *CD69 (Early Activation Marker):* | *AAATCTGTGTCAGTGGATGC* | *TCATTCTTCTCATTCTTGGG* |
| *IFN-gamma* | *GAGTGTGGAGACCATCAAGGAAG* | *TGCTTTGCGTTGGACATTCAAGTC* |
| *IL-2* | *AGAACTCAAACCTCTGGAGGAAG* | *GCTGTCTCATCAGCATATTCACAC* |
| *IL-10* | *GCCTAACATGCTTCGAGATC* | *CTCATGGCTTTGTAGATGCC* |
| *TAP-1* | *GAC AAG AGC CGC TGC TAT TTG G* | *TGATAAGAAGAACCGTCCGAGA* |
| *TAP-2* | *GCC TGT GCT GTT CTC GGG TTC TGC* | *TGTACCAGGTGGGCGTAG* |
| *LMP-2* | *CTC TGC ACC AGC ACA TCT T* | *AGAGTGATGGCATCTGTGGT* |
| *LMP-7* | *ATG GCG TTA CTG GAT CTG TGC GGT GC* | *TCACAGAGCGGCCTC TCCGTACTTGTA* |
| *TIM-3* | *CCAAATCCCAGGCATAAT* | *AAGCGACAACCCAAAGGT* |
| *LAG-3* | *GCAGTGTACTTCACAGAGCTGTC* | *AAGCCAAAGGCTCCAGTCACCA* |

*Table S1: Primers used in RT-PCR studie*

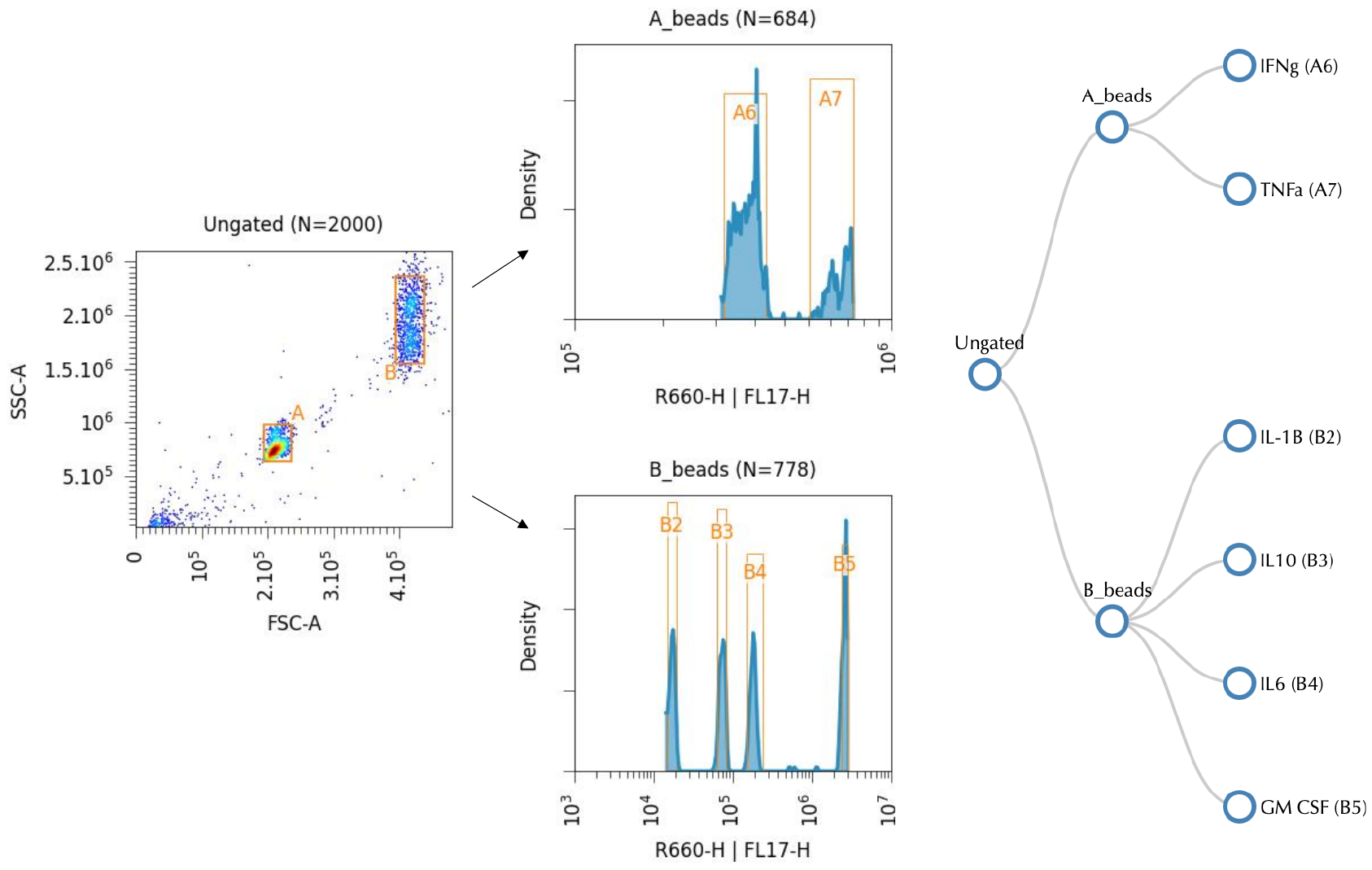

*Figure S7: Gating strategy for Cytokine Bead Array.*

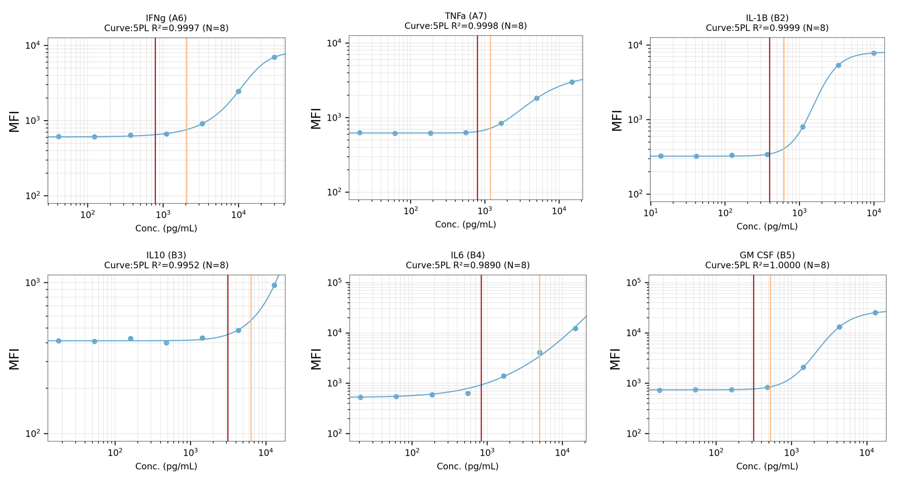

*Figure S8: Standard curves for each cytokine depict R^2^ values above 0.9.*

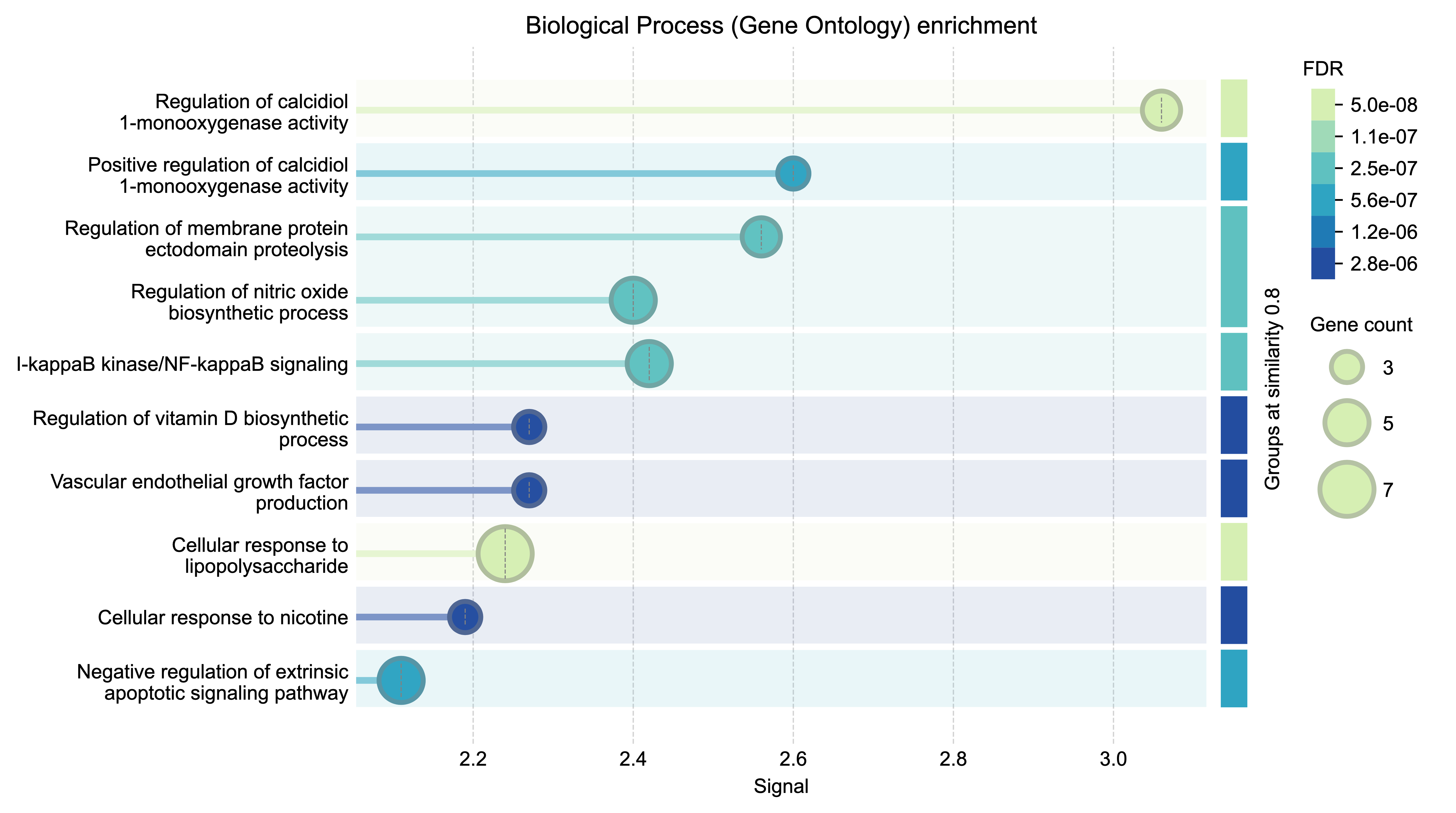

*Figure S9: Functional enrichment visualization for STRING analysis.*
